# Incorporating the image formation process into deep learning improves network performance in deconvolution applications

**DOI:** 10.1101/2022.03.05.483139

**Authors:** Yue Li, Yijun Su, Min Guo, Xiaofei Han, Jiamin Liu, Harshad D. Vishwasrao, Xuesong Li, Ryan Christensen, Titas Sengupta, Mark W. Moyle, Jiji Chen, Ted B. Usdin, Daniel Colón-Ramos, Huafeng Liu, Yicong Wu, Hari Shroff

## Abstract

We present ‘Richardson-Lucy Network’ (RLN), a fast and lightweight deep learning method for 3D fluorescence microscopy deconvolution. RLN combines the traditional Richardson-Lucy iteration with a fully convolutional network structure, improving network interpretability and robustness. Containing only ∼16 thousand parameters, RLN enables 4- to 50-fold faster processing than purely data-driven networks with many more parameters. By visual and quantitative analysis, we show that RLN provides better deconvolution, better generalizability, and fewer artifacts than other networks, especially along the axial dimension. RLN outperforms Richardson-Lucy deconvolution on volumes contaminated with severe out of focus fluorescence or noise and provides 4- to 6-fold faster reconstructions of large, cleared tissue datasets than classic multi-view pipelines. We demonstrate RLN’s performance on cells, tissues, and embryos imaged with widefield-, light sheet-, and structured illumination microscopy.

## Introduction

All fluorescence images are contaminated by blurring and noise, but this degradation can be ameliorated with deconvolution^1, 2, 3^. For example, iterative Richardson-Lucy deconvolution (RLD)^4, 5^ is commonly used in fluorescence microscopy, and is appropriate if the dominant noise source is described by a Poisson distribution. Unfortunately, RLD is computationally taxing for 3D- and 3D timelapse (4D) data, particularly if complex regularization^6, 7^ or large numbers of iterations are applied. To address this challenge, we recently proposed RLD variants^8^ that can accelerate deconvolution speed by at least tenfold by reducing the number of iterations. Deploying these methods requires careful parameter optimization to avoid introducing artifacts.

Parameter tuning is usually experience-dependent and time-consuming, and would ideally be automated. Deep learning offers one route to automation, as neural networks can automatically learn the mapping between the input data and desired output, given ample training data. Many deep learning models now show excellent capability in super-resolution, denoising, and deconvolution applications, including content-aware image restoration networks (CARE)^9^ based on the U-net architecture^10^, residual channel attention networks (RCAN)^11, 12^, and DenseDeconNet (DDN)^8^. Drawbacks of these methods include poor network interpretability and their data-driven nature. The latter implies that the quantity and quality of training data can drastically affect network performance. Another concern with deep learning methods is generalizability, i.e., whether a network trained on one type of data can be used to make predictions on another data type.

Combining the interpretability of traditional model-based algorithms and the powerful learning ability of deep neural networks is a promising approach for avoiding tedious parameter tuning on the one hand and poor generalizability on the other. Algorithm unrolling^13^ provides such a framework, using neural network layers to represent each step in traditional iterative algorithms (e.g. ADMM-net^14^ or ISTA-net^15^). Passing input data through the unrolled network is equivalent to executing the iterative algorithm a finite number of times.

Inspired by RLD and algorithm unrolling, we propose a 3D microscopy deconvolution method that combines the forward/backward projector structure in RL deconvolution and deep learning, i.e., Richardson-Lucy Network (RLN). We benchmarked the deconvolution capability of RLN against traditional RLD and purely-data-driven networks including CARE, RCAN and DDN. We found that RLN causes fewer artifacts than purely data-driven network structures, providing better deconvolution and generalization capability. RLN contains less than one-sixtieth the number of learning parameters than CARE and RCAN, enabling at least 4-fold improvement in processing time. Finally, RLN provides better axial resolution than RLD, even in the low signal-to-noise (SNR) ratio regime and when RLN is trained on synthetic data. We demonstrate the power of RLN on simulated phantoms and diverse samples acquired with widefield-, light-sheet-, and structured illumination microscopy.

## Results

### Richardson-Lucy deconvolution motivates a new network architecture

The update formula in RLD (**Supplementary Fig. 1a, Methods**) can be decomposed into four steps:

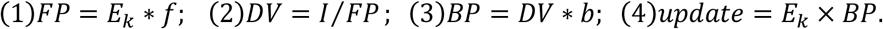

Here the forward projector *FP* function is the convolution of the current object estimate *E*_*k*_ with the forward projector *f, DV* indicates the division of the raw image *I* by *FP*, the backward projector *BP* function is the convolution of *DV* with the backward projector *b*, and the estimate is updated by multiplying *E*_*k*_ with *BP*. With appropriate design of *f* and *b*, the speed of deconvolution can be improved^8^. However, the need to define parameters manually and the challenge of defining a stopping criterion^8^ remain problematic. Given the reliance of RLD and fully convolutional networks on the convolution operation, we wondered whether the latter might be used to find the proper convolution parameters, thereby solving the projector design problem automatically.

We introduced *FP, DV* and *BP* functions into a convolutional network through algorithm unrolling, creating a new 3D microscopy deconvolution network, Richardson-Lucy Network (RLN). The RLN structure consists of three core components: down-scale estimation (H1), original-scale estimation (H2), and merging (H3) (**Fig. 1a, Supplementary Fig. 1b, Methods**). H1 and H2 explicitly follow the RL deconvolution update formula (**Methods**), and H3 merges H1 and H2 with convolutional layers, providing the final deconvolved output.

**Fig. 1.**
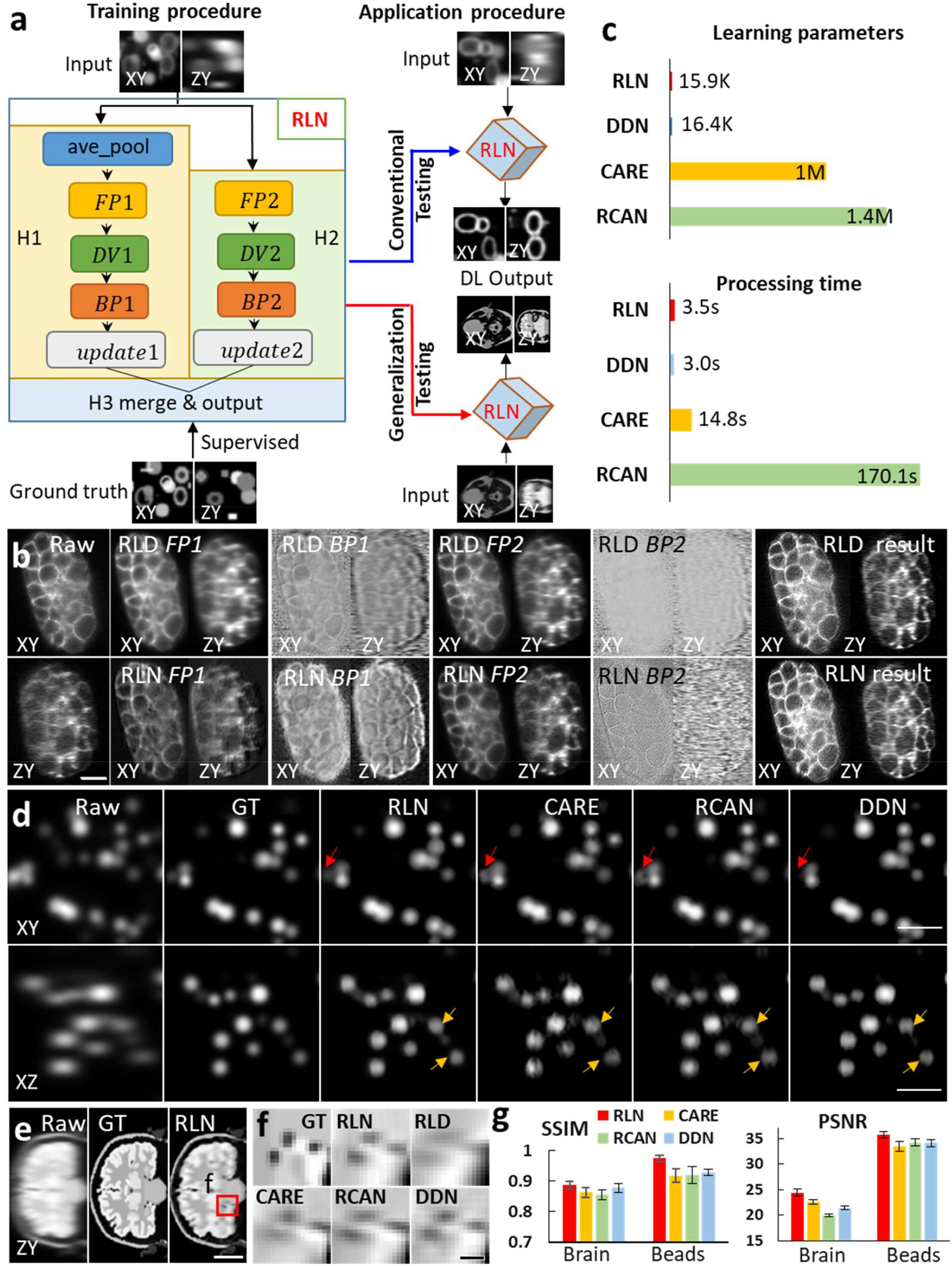
RLN schematic and performance comparison with CARE, RCAN and DDN. **a)** Schematic design of RLN consisting of three parts: H1, H2, and H3. *FP1, DV1, BP1, FP2, DV2, BP2* in H1/H2 follow the RL deconvolution iterative formula. See **Methods** for further details. H3 combines and integrates the feature maps produced by H1 and H2. During training, low-resolution volumes (i.e., input) are fed into RLN, and the corresponding high-resolution ground truth are used to supervise the learning of network parameters. When applying the model, we perform conventional testing (training datasets include the same types of structures as in the test datasets) and generalization testing (i.e., test datasets are from different types of structures than the training datasets). **b)** Example from *C. elegans* embryos expressing GFP-membrane in XY and ZY views of single view data acquired with diSPIM, showing raw input (left column), intermediate outputs (middle columns) and result (right column) of RLN (bottom row) vs. RLD (top row). RLN: *FP1, BP1, FP2, BP2* are the steps in H1 and H2 shown in **Fig. 1a**; RLD: *FP1* and *BP1* are the forward projection and backward projection at iteration 1, and *FP2, BP2* at iteration 5. Similarities between the RLN and RLD intermediates highlight RLN interpretability. **c)** Parameter number and testing runtime for ∼200 MB dataset (1920 × 1550 × 20 voxels), comparing RLN, DDN, CARE and RCAN. RLN offers the fewest parameters and a runtime ∼4-fold faster than CARE and ∼50-fold than RCAN. **d)** Simulated noiseless spherical phantoms in lateral and axial views, comparing raw input, ground truth (GT), and generalization predictions from RLN, CARE, RCAN and DDN. Network predictions are derived from a training dataset with mixed structures consisting of a combination of dots, solid spheres, and ellipsoidal surfaces, emphasizing the generalization capability of RLN. See **Methods**. RLN provides a prediction closest to the ground truth (red and yellow arrows), especially in axial views of the sample. CARE, RCAN and DDN showed distorted shape or information loss (red and yellow arrows). **e)** Simulated 3D noiseless human brain results in axial ZY view. Left to right: raw input, ground truth (noiseless and without blur), and RLN prediction from raw input. **f)** Magnified axial view corresponding to red rectangle in **e)**, showing that RLN provides better restoration than RLD or CARE/RCAN/DDN predictions. See also **Supplementary Fig. 4**. The predictions rely on models trained on the same data as used for **d)**, and thus underscore the generalization capability of RLN. **g)** Quantitative analysis with SSIM and PSNR for the predictions in **d)** and **e)**, confirming that RLN offers the closest match to ground truths. See also **Supplementary Table 2**. Means and standard deviations are reported, obtained from N = 3 subvolumes for brain and 3 volumes for beads. Scale bars: **b)** 10 μm, **d)** 5 μm, **e)** 50 pixels, **f)** 6 pixels.

To demonstrate the interpretability of RLN, we examined the intermediate and final outputs of RLD and RLN on different phantoms and samples (**Supplementary Table 1)**. We studied network performance in conventional tests (where training and test data correspond to the same type of sample) and generalization tests (where the extent to which a model trains on one type of data generalizes to another, **Fig. 1a**). First, we created a synthetic phantom object consisting of mixtures of dots, solid spheres, and ellipsoidal surfaces (**Supplementary Fig. 2**) and evaluated network output on held out data of the same sample type (**Supplementary Fig. 3, Methods**). Second, we evaluated the generalization performance of the mixed structure model when applied to a human brain phantom (**Supplementary Fig. 4a-f, Methods**). Third, we acquired single-view volumes of GFP-labeled cell membranes in a *C. elegans* embryo, acquired with dual-view light-sheet microscopy (diSPIM^16^, **Fig. 1b**) again under conventional testing with single-view input (i.e., the model was trained on similar single-view embryo data). As shown in the lateral and axial views in all these examples, the intermediate output produced by RLN maintain the structure of the input data and resemble the output of RLD. For example, in both RLD and RLN, the *FP* results are blurry, as expected since in both methods this step mimics the blurring introduced by imaging. For the simulated brain phantoms, we found that the final RLN results (structural similarity index^17^ (SSIM) 0.89, peak signal-to-noise ratio (PSNR) 24.4) outperformed RLD (SSIM 0.72, PSNR 16.9) (**Supplementary Fig. 4**), producing reconstructions closer to the ground truth. In all examples, we also noticed that RLN produced sharper axial views than RLD.

To study the effectiveness of the RL structure in RLN, we constructed an ablated version of RLN termed RLN-a (**Supplementary Fig. 1c**) by removing the *DV* and *update* steps. First, we compared intermediate and final outputs from RLD, RLN, and RLN-a on a simulated bead dataset (**Supplementary Fig. 5**), using the model trained on the synthetic mixed structures (**Supplementary Fig. 2**). The intermediate steps in RLN-a appeared visibly different than RLD and RLN (particularly the *FP1* step); and the prediction from RLN-a (SSIM 0.94, PSNR 34.0) was noticeably further from the ground truth than RLN (SSIM 0.97, PSNR 35.7), suggesting that the *DV* and *update* steps in RLN are useful in network generalizability. Next, we compared the deconvolution ability of RLD, RLN and RLN-a in the presence of varying levels of Gaussian and Poisson noise by training models on mixed phantom structures with different input SNR levels (i.e., one model for each SNR level, **Supplementary Fig. 6**). The performance of all methods degraded as the level of noise increased, although RLN performed better than RLN-a, and both networks produced outputs visually and quantitatively closer to the ground truth than RLD at all noise levels. RLD also produced ‘over-deconvolved’ reconstructions relative to RLN and RLN-a. In summary, these results demonstrate the usefulness of the convolutional network structure in RLN-a and RLN in deconvolving noisy data, and that the additional structure in RLN further improves deconvolution output relative to RLN-a.

### Performance of RLN vs. CARE, RCAN and DDN on simulated data

We compared the number of network parameters and the processing time of RLN with other state-of-the-art networks including CARE^9^, RCAN^11^ and DDN^8^ (**Fig. 1c**). Both RLN and DDN are lightweight models, using less than one-sixtieth the number of learning parameters than CARE and RCAN. The time required to train an RLN model is comparable to CARE and DDN, but ∼3x faster than RCAN (e.g., for 100 iterations with 64 × 64 × 64 voxels, RLN required 26.5 s vs. 29.2 s with CARE, 23.6 s with DDN, and 90.9 s with RCAN). When applying the models to deconvolve sample volumes of ∼200 MB size (1920 × 1550 × 20 voxels), RLN and DDN required ∼3 s vs. ∼15 s with CARE and ∼170 s with RCAN.

Next, we simulated noiseless spherical phantom datasets to examine (1) the difference in RLN reconstructions in conventional testing vs generalization applications (**Supplementary Fig. 7)**; and (2) the output of RLN vs. CARE, CAN and DDN (**Fig. 1d)**. Ground truth spherical structures were generated by ImgLib2^18^ and blurred with a Gaussian kernel, and input data were generated by further blurring ground truth structures with the point spread function (PSF) of the 0.8/0.8 NA diSPIM (**Methods**). Generalization tests were conducted using the models trained from the synthetic mixed structures. RLN under both conventional testing and generalization paradigms recovered axial views distorted by the PSF and provided better linearity than RLD (**Supplementary Fig. 7)**, although the generalized prediction was artificially sharpened compared to the conventional result. Notably, RLN offered better generalization than CARE, RCAN and DDN, all of which had more obvious visual distortions (red and yellow arrows, **Fig. 1d**) and lower SSIM and PSNR (**Fig. 1g**).

Last, we applied the models trained from the synthetic mixed structures (**Supplementary Fig. 2**) to the synthetic human brain phantom (**Fig. 1e, Supplementary Fig. 4g**), which is visually very different than the structures in the training data. RLN and DDN generalization outputs more closely resembled the ground truth than RLD or CARE/RCAN generalization outputs (axial views shown in **Fig. 1f**, and lateral views shown in **Supplementary Fig. 4g**), a result consistent with SSIM and PSNR analysis (**Fig. 1g, Supplementary Table 2**).

### RLN improves deconvolution capability relative to other networks on biological images

To demonstrate the deblurring capability of RLN for biological images, we used previously published images of live U2OS cells transfected with mEmerald-Tomm20, acquired with dual-view light sheet fluorescence microscopy (0.8/0.8 NA diSPIM)^8, 19, 20^. Here we trained RLN, CARE, RCAN and DDN models to predict the dual-view, joint deconvolved results based on 12 randomly selected volumes from the time series. The training pairs consisted of raw single view data paired with corresponding dual-view joint deconvolution ground truth (**Supplementary Tables 3, 4** provide additional information on the training parameters used in each model). RLN prediction showed clear improvements in resolution and contrast compared to the raw input (**Fig. 2a**), especially in the axial direction. Compared with other networks, mitochondrial details revealed by RLN were more similar to the ground truth in both lateral (**Fig. 2b**) and axial views (**Fig. 2c**), both visually and via quantitative assessment (**Fig. 2f**). We observed similar improvements when applying the RLN model to another live U2OS cell transfected with mEmerald-Tomm20, acquired with 0.8/0.8 NA diSPIM every 3 seconds, over 200 time points **(Supplementary Video 1**).

**Fig. 2.**
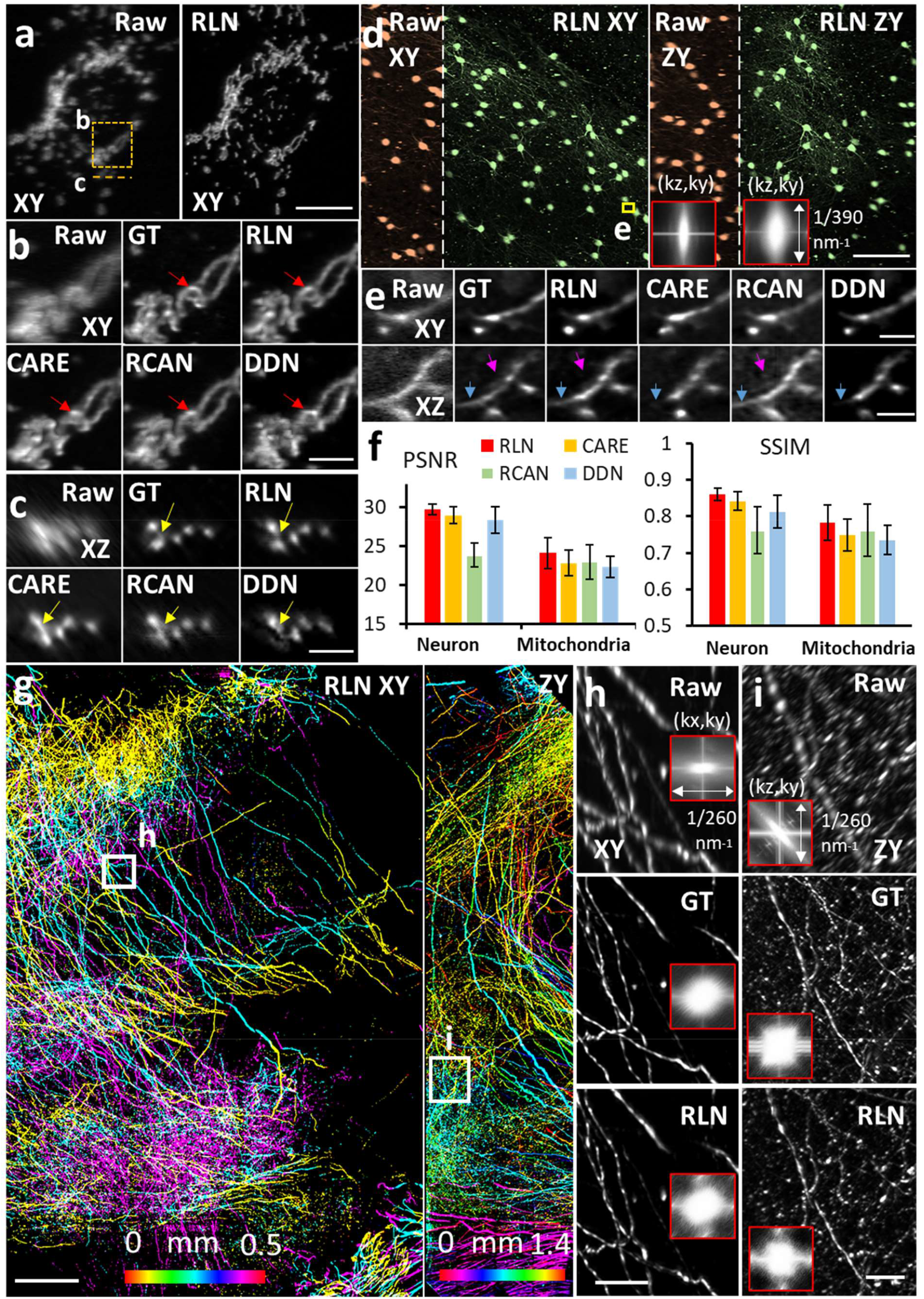
Deconvolution ability of RLN on thin or cleared biological samples. **a)** Live U2OS cells transfected with mEmerald-Tomm20 were imaged with symmetric 0.8/0.8 NA diSPIM. Lateral maximum intensity projections (left: raw single view; right: RLN prediction, conventional testing with single-view/joint deconvolution training pairs) are shown for a single time point. **b, c)** Higher magnification of dashed orange line, rectangle in **a)**, highlighting fine mitochondrial features (circular shape, red arrows; separated mitochondrial cross sections, yellow arrows) in lateral **b)** and axial view **c)**, comparing raw single-view input, dual-view joint deconvolution ground truth, predictions from RLN, CARE, RCAN, and DDN. Visually, the RLN output most closely resembles the ground truth. **d)** XY and ZY maximum intensity projections of cleared brain tissue slab expressing Alexa Fluor 555-conjugated secondary antibody against anti-tdTomato primary antibody, acquired by cleared-tissue immersion 0.4/0.4 NA diSPIM. This volume spans 1184 × 1184 × 1218 voxels, stitched from 25 subvolumes, each with size 288 × 288 × 1218 voxels. Orange: raw data, green: RLN prediction. The inset panel in the ZY view shows the Fourier spectra of raw input and RLN output, indicating improvement in resolution after RLN. **e)** Higher magnification of yellow rectangle in **d)**, highlighting fine neurites in lateral and axial view, comparing raw, dual-view deconvolution ground truth, predictions from RLN, CARE, RCAN, and DDN. As shown, RLN provides the most similar results to the ground truth while other methods show information loss or blurry details. **f)** Quantitative analysis with SSIM and PSNR for data shown in **a)** and **d)**. See also **Supplementary Table 2**. Means and standard deviations are reported, obtained from N = 3 subvolumes within the large neuron volume and 3 volumes from time-lapse mitochondria data. **g)** Lateral (left) and axial (right) maximum intensity projections from a cleared brain tissue slab expressing tdTomato in axons, acquired with 0.7/0.7 NA cleared tissue diSPIM and recovered with RLN. This volume spans 5432 × 8816 × 1886 voxels (∼1.4 × 2.3 × 0.5 mm^3^), and is stitched from 900 subvolumes each with size 1500 × 1500 × 42 voxels. Images are depth coded as indicated. **h, i)** Higher magnification views of lateral (**h**) and axial (**i**) projections corresponding to white rectangular regions in **g)**, comparing fine neurites and corresponding resolution estimates (Fourier spectra shown in the inserts) as assessed with raw single view (top), dual-view deconvolved ground truth (middle), and RLN prediction (bottom). The RLN output closely resembles the ground truth with an SSIM of 0.97 ± 0.03, PSNR 49.7 ± 2.2 (N = 80 XY slices). Scale bars: **a, e)** 10 μm, **b, c)** 5 μm, **d)** 100 μm, **g)** 200 μm, **h, i)** 50 μm.

Next, we examined images of neurites in a slab of mouse brain, acquired by cleared-tissue diSPIM with 0.4/0.4 NA lenses^8^. The mouse brain sample was prepared using the iDISCO+ procedure^21^, followed by immunolabelling with Alexa Fluor 555-conjugated secondary antibody against anti-tdTomato primary antibody. The entire brain volume after dual-view reconstruction spanned 10280 × 5160 × 1400 voxels (corresponding to 4 × 2 × 0.5 mm^3^, 138.3 GB in 16-bit format)^22^. We trained on 12 randomly selected subvolumes of single-view data to predict the dual-view joint deconvolved results, each comprising 128 × 128 × 128 voxels, then applied the model to a held-out, larger scale subvolume from the same dataset spanning 1184 × 1184 × 1218 voxels. RLN provided the best visual output of the neurites in both lateral and axial views (**Fig. 2d**), compared to RCAN, CARE and DDN (**Fig. 2e**), again confirmed quantitatively via PSNR and SSIM (**Fig. 2f, Supplementary Table 2**). In this example, we cropped the larger subvolume into 25 batches, processed each subvolume with RLN, and stitched the deep learning output to generate the final reconstruction (**Methods**). Cropping, RLN prediction, and stitching took ∼3 minutes. Scaling up this RLN processing routine to the whole brain slab implies a time of ∼2.2 hours with the RLN pipeline, a 5.5-fold speed up compared to the conventional processing pipeline described in our previous publication^8^, which would otherwise take 12 hours including cropping, registration, joint RL deconvolution, and stitching.

To further demonstrate that RLN can accelerate the restoration of large datasets, we imaged a large multi-tile image volume from another slab of cleared mouse brain, this time at higher resolution with a 0.7/0.7 NA cleared tissue diSPIM (**Methods**). This brain expressed tdTomato in axonal projections from the site of a stereotactic AAV injection. After fixing, clearing, and sectioning the brain, we located and imaged a region with dense neurite labelling. The size of the brain volume after dual-view reconstruction spanned 5432 × 8816 × 1886 voxels (∼1.4 × 2.3 × 0.5 mm^3^, 168.2 GB in 16-bit format). From this dataset we randomly selected 40 subvolumes, each 256 × 256 × 256 voxels, pairing single-views (input) with dual-view joint deconvolutions (ground truth) to train the network, then applied the trained model to the entire dataset (**Fig. 2g, Supplementary Video 2)**. Cropping the entire volume into 900 subvolumes, performing the RLN prediction on each subvolume, and stitching the results back together took ∼2.7 hours on a single workstation equipped with a consumer-grade GPU card. Compared to the single-view raw data, the RLN prediction displayed improved image resolution and contrast, closely resembling the joint deconvolution ground truth in both lateral (**Fig. 2h)** and axial (**Fig. 2i)** views (SSIM of 0.97 ± 0.03, N = 80 XY planes). By contrast, it took approximately 11.5 hours to run the registration, joint deconvolution and stitching with the same GPU card as used above for applying RLN, or 3.5 hours on a cluster (**Methods**). We further tested the speed-up of RLN processing time with different sizes of data (∼3-300 GB), confirming that RLN provides a 4-6-fold speed improvement (**Supplementary Fig. 8a**) over the previous processing pipeline^8^ for cleared-tissue diSPIM data restoration. Finally, we note that the registration necessary for dual-view fusion failed on a small number of subvolumes in this example (e.g., with sparse signals), causing artifacts in the joint deconvolution result. As RLN is applied only on single-view input, it completely avoids errors of this kind (**Supplementary Fig. 8b, c)**.

Although the RLN prediction with single-view input (in the relatively thin or transparent samples studied thus far) closely resembled the dual-view ground truth, fine axial detail present in the ground truth was not fully recovered (e.g., compare ground truth to RLN predictions in axial views, **Fig. 2c, e, i**). To address this issue and generate more nearly isotropic reconstructions, particularly in the context of densely labeled and scattering samples where additional views provide critical information lacking in any single view, we developed a dual-input implementation of RLN (**Methods, Supplementary Fig. 9**). We then tested this modified RLN with two inputs corresponding to the two raw registered views acquired with diSPIM. After training the dual-input RLN model with 12 registered dual-view volumes acquired by imaging living GFP-histone-labeled *C. elegans* embryos with a 0.8/0.8 NA diSPIM, applying the model produced better reconstructions than did the single-view RLN (**Supplementary Fig. 10a, b, g, Supplementary Video 3**). We observed similar improvements on embryos labeled with GFP-membrane markers (**Supplementary Fig. 10c-g**).

### RLN generalizes well on biological data

Having demonstrated RLN generalizability on simulated data (**Fig. 1d, e, Supplementary Fig. 4**), we next turned to biological data. We found that training models on simulated mixed structures that were blurred with the diSPIM PSF (**Methods**) generalized well when applied to *C. elegans* embryos labeled with nuclear and membrane markers and imaged with diSPIM (**Supplementary Fig. 11**). Although the generalization result was slightly inferior to the conventional test result (i.e., training directly on diSPIM data), it still compared favorably against single-view RLD.

Next, we examined RLN generalizability on super-resolution data. We began by imaging (1) mitochondria labeled by mEmerald-Tomm20-C-10 (Mito) and (2) the endoplasmic reticulum (ER) labeled by ERmoxGFP in live U2OS cells with iSIM^23^, a rapid super-resolution microscopy technique. We trained two RLN models with Mito and ER training data, respectively, and performed cross-validation testing: (1) the Mito data were predicted with the model trained with Mito and with the model trained with ER (**Fig. 3a, b**); (2) the ER data were similarly predicted by the two models (**Fig. 3c**). Similar to the deconvolved ground truth, RLN enabled crisper visualization of Mito and ER compared to the raw input. The predictions from different RLN models were nearly identical, both with SSIM higher than 0.96 and PSNR higher than 39 dB (**Fig. 3d**). These results suggest that the RLN predictions based on super-resolution input do not rely exclusively on image content, implying that gathering ground truth data on a single type of structure is likely sufficient to predict another type of structure. Further cross-testing experiments on lysosomal and Golgi markers support this claim (**Supplementary Fig. 12**).

**Fig. 3.**
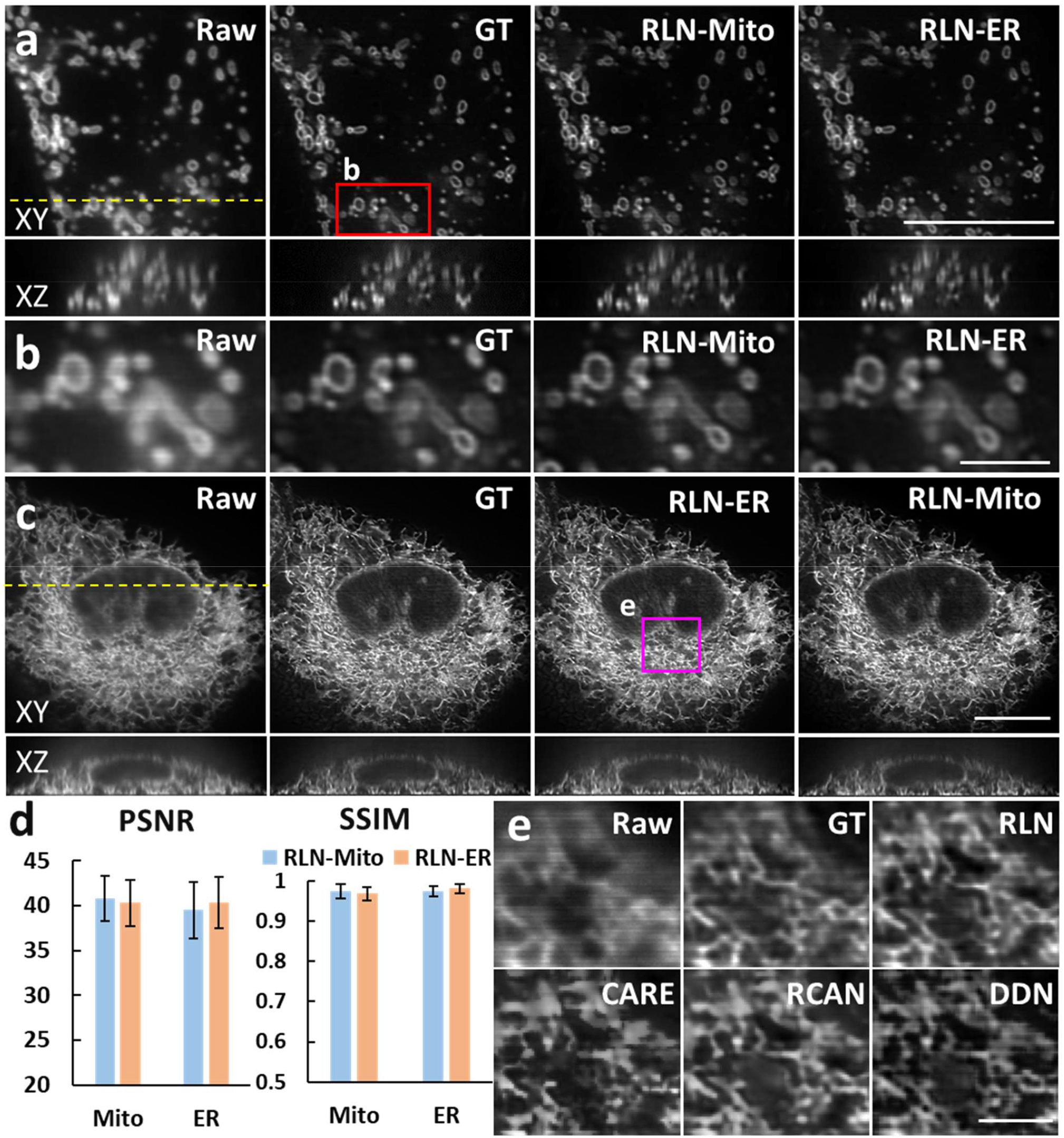
RLN generalizes well on biological samples. **a)** Lateral (top) and axial (bottom) views of live U2OS cells expressing mEmerald-Tomm20-C-10, acquired with iSIM, comparing the raw input (i.e., without deconvolution), ground truth (the RL deconvolved result), the prediction from Mito-trained RLN (RLN-Mito) and the prediction from ER-trained RLN (RLN-ER). Axial views are taken across yellow dashed line. **b)** Higher magnification of red rectangle in **a)**, further highlighting fine mitochondrial details. **c)** Lateral and axial views of live U2OS cells expressing ERmoxGFP, acquired with iSIM, comparing raw input, ground truth, predictions from Mito-trained RLN (RLN-Mito) and ER-trained RLN (RLN-ER). Axial views are taken across the yellow dashed line. **d)** Quantitative analysis with SSIM and PSNR for **a-c)**, indicating the good generalizability of RLN. Means and standard deviations are obtained from N = 6 volumes for Mito and ER. **e)** Magnified view of the magenta rectangle in **c**), comparing raw, RL deconvolved ground truth, and predictions from RLN, CARE, RCAN, and DDN. See **Supplementary Table 2** for the quantitative SSIM and PSNR analysis of these network outputs. Note that for data shown in **e)**, all network models were trained with a simulated phantom consisting of dots, solid spheres, and ellipsoidal surfaces. See **Supplementary Figs. 12, 13** for additional comparisons on other organelles. Scale bars: **a, c, f)** 10 μm, **b, e)** 2 μm.

We then compared the generalization ability of RLN to other deep learning models (CARE, RCAN and DDN) on biological data. First, we found that RLN provides better deconvolution on Mito data than DDN, CARE and RCAN (**Supplementary Fig. 13a-c, Supplementary Table 2)**, when using models trained on ER. Second, we compared the output of RLN, CARE, RCAN, and DDN models trained exclusively on the synthetic mixed structures (**Supplementary Fig. 2**, blurred with the iSIM PSF^11^, **Methods**). When applying such models to the ER and lysosome biological data, we again found that RLN gave superior results, showing fewer artifacts and more closely resembling the ground truth than other networks (**Fig. 3e, Supplementary Fig. 13d-f, Supplementary Table 2**).

### RLN outperforms direct RLD on volumes contaminated by severe background or noise

Deconvolving volumes that are badly contaminated with background or noise is challenging. To illustrate the potential of RLN to address the former, we examined two samples. First, we used widefield microscopy to image fixed U2OS cells stained in four colors for actin, tubulin, mitochondria, and nuclei (**Methods, Fig. 4a)**. Then we trained an RLN model with synthetic mixed structures (**Supplementary Fig. 2**) and applied the model to these widefield data. RLN outperformed RLD in lateral and axial views, sharpening nuclei, resolving more mitochondria, better separating actin and microtubule filaments, and recovering high spatial frequencies otherwise swamped by background (insets in **Fig. 4b-c, Supplementary Fig. 14**). We also applied the RLN model trained with simulated data to a *C. elegans* embryo expressing ttx-3B-GFP, imaged with widefield microscopy (**Methods, Fig. 4d)**. This marker labels neuronal membranes in the animal’s head, and leaky expression from unc-54 3’UTR likely labels membranes in gut cells. Compared to RLD, RLN better distinguished neuronal cell bodies and two functionally distinct nerve bundles - the sublateral and sensory neuron bundles within the nerve ring (main neuropil of *C. elegans*), which are challenging to distinguish due to the small size of the embryonic nerve ring (**Fig. 4e-f)**. RLN also better restored gut cell membranes than RLD (**Supplementary Fig. 15)**.

**Fig. 4.**
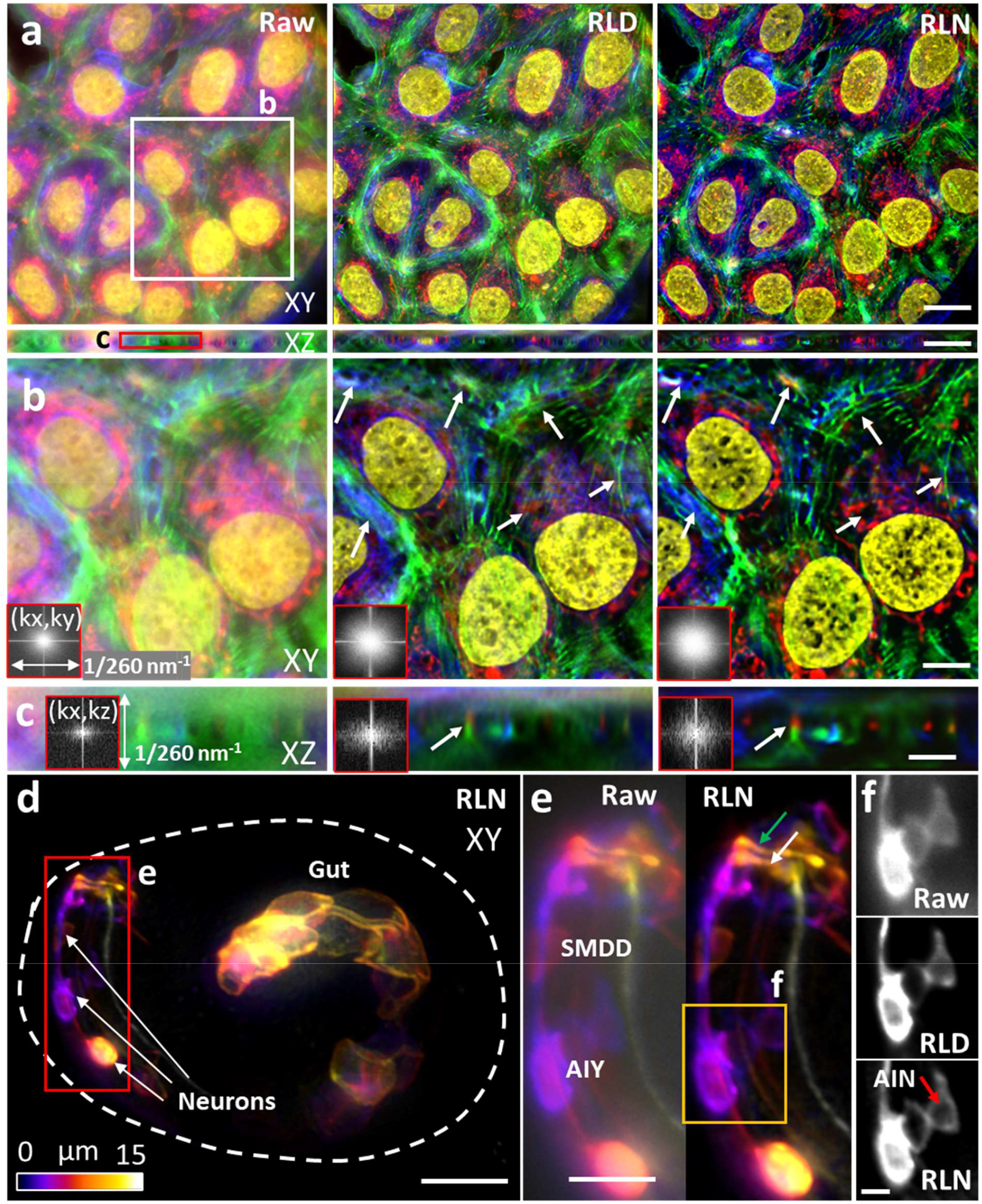
RLN trained with synthetic mixed structures outperforms direct RLD on volumes contaminated by severe out-of-focus background. **a)** Four color lateral and axial maximum intensity projections of a fixed U2OS cell, acquired by widefield microscopy, comparing the raw input, RLD, and RLN prediction based on a model trained on synthetic mixed structures (**Supplementary Fig. 2**). Red: mitochondria immunolabeled with anti-Tomm20 primary antibody and donkey α-rabbit-Alexa-488 secondary; green: actin stained with phalloidin-Alexa Fluor 647; Blue: tubulin immunolabeled with mouse-α-Tubulin primary and goat α-mouse-Alexa-568 secondary; yellow: nuclei stained with DAPI. **b, c)** Higher magnification views of white and red rectangular regions in **a)** at a single slice, highlighting fine structures (white arrows) that are better resolved with RLN prediction than RLD in lateral **b)** and axial views **c)**. Fourier spectra of the sum of all channels shown in the inserts also indicate that RLD better recover resolution than RLD. See **Supplementary Fig. 14** for the Fourier spectra of individual channels. **d)** Depth-coded image of a *C. elegans* embryo expressing ttx-3B-GFP, acquired by widefield microscopy, and predicted by RLN based on a model trained on the synthetic mixed structures. Dashed line indicates the embryo boundary. **e)** Higher magnification of red rectangle in **d)**, comparing the raw input and RLN prediction, showing neuronal cell bodies (AIY and SMDD) and neurites (the sublateral neuron bundle, green arrow; the amphid sensory neuron bundle, white arrow) are better resolved with RLN. **f)** Higher magnification of orange rectangle in **e)**, comparing the raw input, RLD, and RLN prediction, highlighting AIY and AIN neurons. Red arrow highlights interior of neuron, void of membrane signal and best resolved with RLN. See **Supplementary Fig. 15** for images showing comparisons of gut cell membranes. Scale bars: **a)** 20 μm, **b, d)** 10 μm, **c, e)** 5 μm, **f)** 2 μm.

Another class of problematic samples for conventional RLD concerns those with poor SNR. RLN prediction is also influenced by the SNR of the input data. In addition to investigating the noise dependence of RLN on simulated data **(Supplementary Fig. 6)**, we also studied biological samples, imaging U2OS cells expressing ERmoxGFP with instant SIM^23^. When we trained an RLN model to deconvolve noisy data (input with an SNR of ∼5, ground truth with SNR of ∼40), the prediction was visually improved compared to either the raw input or the RLD result, which were each dominated by noise (**Supplementary Fig. 16**). Considering that using two or more networks sequentially can provide better restoration than a single network^11, 24^, we also performed a two-step deep learning strategy, first applying a denoising RCAN model to initially improve SNR, then applying a deconvolution RLN model for further improvement of contrast and resolution. In this two-step training scheme, the first step RCAN model was trained on pairs of low/high SNR raw data, whereas the second step RLN model was based on pairs of high SNR raw data and high SNR deconvolved data. We found the two-step prediction was noticeably closer to the ground truth and provided higher PSNR and SSIM than the single-step prediction with either RCAN or RLN model alone (**Supplementary Fig. 16**).

## Discussion

We designed RLN to mimic the forward/backward projector architecture of classic iterative deconvolution (**Fig. 1a, Supplementary Fig. 1**), thereby improving network interpretability and performance (**Fig. 1b, Supplementary Figs. 3-5**). Since parameters are learned automatically, RLN has the potential to eliminate manual parameter selection in state-of-the-art deconvolution, as well as the burdensome and currently unsolved stopping criterion problem^8^. RLN is designed with ∼16,000 parameters, ∼60-90 fold fewer than purely-data-driven network structures like RCAN and CARE (**Fig. 1c**). RLN also offers rapid run time after training, more than 4-fold faster than CARE, and almost 50-fold faster than 3D RCAN (**Fig. 1c**). With this advantage, it offers 4-6-fold speed up (**Supplementary Fig. 8a**) over our previous processing pipeline for the reconstruction of large, cleared tissue datasets with dual-view light sheet microscopy^8^ (**Fig. 2d, e, g-i**). Because the single-view RLN prediction displayed improved resolution and contrast against the raw input, closely resembling the joint deconvolution ground truth (**Fig. 2**), it can reduce the total amount of data required, and bypass artifacts induced by poor registration (**Supplementary Fig. 8b, c)**.

Compared with purely-data-driven network structures, RLN shows better performance on both simulated data (**Fig. 1d, e, Supplementary Fig. 4**) and biological data derived from light-sheet (**Fig. 2a-f**) and super-resolution microscopes (**Fig. 3e, Supplementary Fig. 13**). As expected, the deconvolution performance of RLN deteriorates in the presence of increasing noise, although RLN still outperforms RLD in the low SNR regime (**Supplementary Figs. 6, 16**). Although it is not always possible to generate high SNR, high quality deconvolved ground truth, the excellent generalization capability offered by RLN suggests that in these difficult cases it may be possible to use synthetic data (**Supplementary Fig. 2**) for training RLN and applying the model to the biological samples (**Figs. 3e, 4, Supplementary Figs. 11, 13d-f**).

We envision several extensions to our work. First, since we have shown that we can successfully use synthetic data to train RLN, our method has the potential to aid in the deconvolution of any multi-view microscope system if the PSF can be defined (e.g., as in our recently published multi-view confocal super-resolution microscopy method^24^). Second, since synthesizing data blurred with different PSFs is trivial, it would be interesting to explore whether RLN can be used to recover images that suffer from spatially varying aberrations or blurring^8,25^. Last, RLN may offer speed or performance improvements over RLD and previous deep learning methods that have been used to reconstruct images acquired with light-field microscopy^26, 27^.

## Data and code availability

The data that support the findings of this study are available upon request. Code for the simulation of 3D mixture phantoms, generation of simulated input data and RLN training/prediction are available at https://github.com/MeatyPlus/Richardson-Lucy-Net

## Author Contributions

Conceived the project: Y. W., H. L. Developed and tested deep learning algorithms/software: Y. L., Y. W. Designed and performed experiments: Y. W., Y. S., X. H., H. V., J. C., H. S. Built instrumentation: Y. W., M. G., H. V. Contributed reagents and advice on biological questions and interpretations: T. S., M. W. M., T. B. U., D. C. R. Processed and analyzed data: Y. L., Y. S., M. G., J. L., X.L., T. S., R.C., Y.W., H.S. Wrote manuscript: Y. L., Y. W., H. S. Supervised research: H. S.

## Acknowledgements

This research was supported by the intramural research programs of the National Institute of Biomedical Imaging and Bioengineering, National Institutes of Health. Y. L. and H. L. acknowledge support from the National Key Research and Development Program of China (2020AAA0109502), the National Natural Science Foundation of China (U1809204, 61701436) and the Talent Program of Zhejiang Province (2021R51004). T. B. U. acknowledges support from the intramural research programs of National Institute of Mental Health (NIMH), National Institutes of Health (ZIC MH002963-05). We thank the Marine Biological Laboratories (MBL), for providing meeting and brainstorming platform. H. S. and D. A. C-R. acknowledge the Whitman and Fellows program at MBL for providing funding and space for discussions valuable to this work. Research in the D. A. C-R lab was supported by NIH grant No. R24-OD016474. M. W. M. was supported by NIH by F32-NS098616. We thank Henry Eden and Patrick La Riviere for careful read and valuable feedback on the manuscript. We thank Zhirong Bao for providing the OD58 *C. elegans* strain used in Fig. 1b and Supplementary Figs. 10-11, Johnny Bui, Grant Kroeschell, and Matthew Chaw for maintaining the worm strains, and W. S. Young (NIMH) for providing early access to the V1b mouse line. This work used the computational resources of the NIH HPC Biowulf cluster (http://hpc.nih.gov).

## Disclaimer

The NIH and its staff do not recommend or endorse any company, product, or service.

## Methods

### Richardson-Lucy network

RL deconvolution (Equation 1) has a compact update structure, needing only one formula to update each estimate:

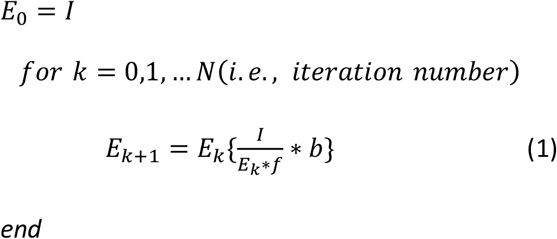

where *I* and *E*_*k*_ are the raw input and estimate of the *k*-th iteration and *f* and *b* are the forward projector (system PSF) and backward projector, respectively. Traditionally *b* is taken to be the transpose of *f*, but using unmatched back projectors (e.g., Gaussian, Butterworth, or Wiener-Butterworth filters)^8^ can result in faster deconvolution by reducing the total number of iterations *N* needed for achieving a resolution limited result.

The key procedure in RLD is convolution. Similarly, convolutional layers are integral to the architecture of deep learning networks, which can learn the convolution kernels automatically. This similarity inspired us to think of using convolutional layers to mimic the convolution with PSF kernels in RL deconvolution. RLN can be regarded as an algorithm unrolling method which uses convolutional layers in a fully convolutional network to represent the convolution steps in each RLD iteration, thereby mimicking the forward/back projection steps.

RLN consists of three parts: H1, H2, H3 (**Fig. 1a, Supplementary Fig. 1b**). H1 functions similarly to an early iteration in RLD, providing a rough estimate of the final output; H2 acts as a late iteration, using all the information in *I* to refine the rough estimate; and H3 is used to merge and integrate the information provided by H1 and H2. The architecture of H1 and H2 closely follow the RL deconvolution update formula, i.e., they mimic the FP and BP steps with convolutional layers, additionally incorporating the division 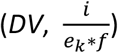, and *update* steps to learn the correction necessary for improving *I*. In RLD, FP and BP procedures use relatively large PSF kernels (e.g., 128 × 128 × 128 voxels for 0.8 NA/0.8 NA diSPIM). Applying such large kernels in a neural network would degrade training efficiency. Typically, deep learning networks use small convolutional kernels with several convolutional layers to extract features. For efficient operation, larger convolution kernels can be replaced by several smaller convolution kernels^28^, e.g. a layer of 5 × 5 convolutions can be replaced by two layers each with 3 × 3 convolutions. To maintain network efficiency, H1 uses smaller feature maps and more layers, while H2 uses larger feature maps and less layers.

Because H1 only roughly estimates the ground truth, it starts with an average pooling layer to down-scale the input volume (i.e., the normalized microscope acquisition *I*) by two in all dimensions to obtain *I*_*ap*_ (average-pooled input). Although this step may cause information loss, it has the benefit of increasing the field of view (FOV), including more spatial information around each voxel, and decreasing computational cost. Following the RL iteration update process, *I*_*ap*_ passes through three convolutional layers to construct the forward projection step. We use dense connections^29^ among these convolution layers, i.e., the outputs of the first two layers are concatenated along the channel direction to act as the input of the third layer, for efficient use of the feature maps. There is also a residual connection between the output feature maps of the third convolution layer and *I*_*ap*_, and the result of this residual connection is denoted *FP1*. This residual connection has two functions: (1) the output of the forward projector *FP* in RL deconvolution is a blurry copy of the current estimate, which approximates the microscope acquisition; and the residual connection acts similarly, adding information learned by the network to the current estimate *I*_*ap*_; (2) it avoids the risk of dividing by zero in the following division step, which may introduce instability in training. All channels of the residual connection are merged by a channel-wise average (C_AVE) producing *FP*1, and the quotient is computed as *DV*1 = *I*_*ap*_/*FP*1. For the back projection step, RLN uses *DV1* as the input to three densely connected convolutional layers to construct *BP1*. Because the final feature maps of H1 need to be restored to the original size, BP1 is up-scaled by a combination of an up-sampling layer and a convolutional layer to obtain the up-scaled *BP1* (*BP1up*). All channels of *BP1up* are merged by a channel-wise average to obtain the correction, which is multiplied with *I* to obtain the estimate *E1*.

H2 is constructed similarly to H1. The differences are that in the *FP* and *BP* steps, there are only two convolutional layers without dense connections; the input of H2 is the original-scale input *I*, i.e., there is no up-scaling procedure; and the correction is applied to *E1* to compute the second estimate *E2*. Since H1 already produces a rough estimate, H2 can use fewer parameters. We thus decreased the number of convolutional layers in H2 to improve memory efficiency.

H3 consists of three convolution layers and uses dense connections to merge and fine-tune *E1* and *E2*. After the channel-wise average of the last layer’s feature map, we obtain the final output *O*.

All convolution layers use [3 × 3 × 3] kernels with [1 × 1 × 1] strides, and are followed with batch normalizations (BN)^30^ and softplus nonlinear activation functions (SP)^31^. The up-sampling consists of transpose convolution operations using [2 × 2 × 2] kernels with [2 × 2 × 2] stride, followed with BN and SP. The SP activation function is a smooth ‘ReLU’ function which ensures non-negativity and avoids ‘dead regions’ where the gradient vanishes and parameters never update. For *DV1* and *DV2*, we add a small constant α = 0.001 in the denominator to prevent division by zero. In the unmatched forward/back projectors design^8^, the choice of forward projector is set to the system PSF while the design of the back projector is more flexible, and should take noise amplification into account. Given that the design of the back projector is more complex, we set the number of output channels of the convolutional layers in the forward projector to 4 and in the backwards projector to 8 to place more weight on learning the back projectors. The total number of parameters in the RLN is ∼16 thousand. For dual-view input, the dual-view information is registered with the ImageJ plugin diSPIM Fusion^8^. RLN merges these registered views by averaging prior to applying H1 (**Supplementary Fig. 9**).

To verify the effectiveness of the *DV* steps and update steps in RLN, we constructed an ablated version of RLN, named RLN-a (**Supplementary Fig. 1c**). RLN-a has the same convolutional layer design as the RLN but removes the *DV* and *update* steps. It shares the same loss function and training parameters as RLN.

In the training procedure, the loss function is given by:

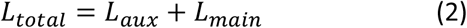

where *L*_*aux*_ is an auxiliary loss term used to guide H1 training, and *L*_*main*_ is the main loss term used to guide training of the whole network.

As *E1* is the rough estimate of the ground truth, it is expected to be sharper than the input volume *I* but blurrier than the ground truth *GT*. Thus, we define intermediate *ITM*:

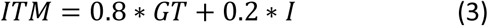

and compute *L*_*aux*_ as the mean square error (*MSE*) between *E1* and *ITM*:

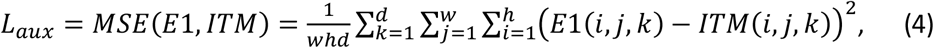

where d, w, h represents the depth, width, and height of the ground truth, respectively.

*L*_*main*_ includes two parts: the mean square error (*MSE*) and structural similarity index^17^ (*SSIM*) between the network output *O* and *GT*:

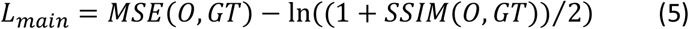

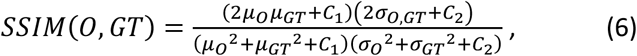

where *μ*_*GT*_, *μ*_*O*_ are the mean values of the *GT* and *O*; 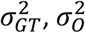 are the variances of the *GT* and *O*; *σ*_*GT*_,_*O*_ is the covariance of *GT* and *O*; and *C*_1_ and *C*_2_ are small constants that prevent the denominator from becoming zero (here *C*_1_ = 1*e*^−4^ and *C*_2_= 9*e*^−4^). A higher *SSIM* value means the network output is more similar to the ground truth. Because the *SSIM* value is smaller than 1, the ln(·) operation is used to keep the loss positive. The *MSE* term is similar to *L*_*aux*_, ensuring that the difference between network outputs and ground truth is as small as possible, but using MSE exclusively may lead to blurred output. *SSIM* is used to preserve the global structural similarity between *O* and *GT*.

The solver method which is used to guide the parameter update is based on the ‘adaptive moment estimation (Adam)’ algorithm. The learning rate *r* decays during the training procedure according to:

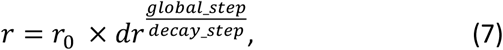

where *r*_*0*_ is the start learning rate, *dr* is the decay rate, *global*_*step* represents the number of training iterations (updated after each iteration), and *decay*_*step* determines the decay period.

Gaussian filter kernels are used to initialize the convolutional layers in *FP*, which contain 4 output channels. Each channel is a Gaussian filter with standard deviation σ = 0.5, 1, 1.5, 2, respectively:

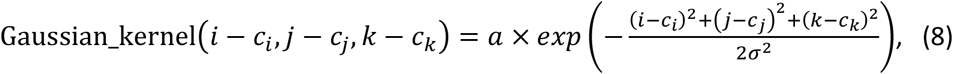

where (*c*_*i*_, *c*_*j*_, *c*_*k*_) is the center coordinate of the kernel and *a* is a random number to increase randomness (ranges from 0.5 to 1). Other kernels in the convolutional layers are randomly initialized with a Gaussian distribution (mean = 0, standard deviation = 1). Using our workstation (see below for details), training with 200 epochs usually takes 2-4 hours, with each epoch using 100 iterations.

Real microscopy volumes often exhibit isolated voxels with bright values that represent abnormal structures. Therefore, we adopted the percentile-based normalization as in CARE^9^:

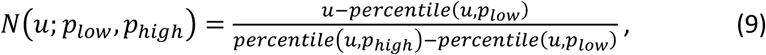

where *percentile*(*u, p*) is the *p*-th percentile of all voxel values of data *u*. For real data, we set *p*_*low*_ ∈ (0,1) and *p*_high_ ∈ (99.0, 100) according to the data quality. For simulated data, we set *p*_*low*_ = 0 and *p*_*high*_ = 100.

We adopted similar online data augmentation as used with 3D RCAN^11^, which is a stochastic block selection process. For every training iteration, the batch size is set to 4. The parameters of RLN and the size of selected blocks are summarized in **Supplementary Table 3**. For the comparison of RLN with RLD, we implemented both conventional RLD (**Figs. 1b, 1f, 4, Supplementary Figs. 3e, 4-6, 11, 14-16**) and RLD with an unmatched back projector (**Fig. 2e, h, Supplementary Fig. 8b, c**). Iteration numbers are included in **Supplementary Table 3**.

### RLN comparison with CARE, RCAN and DDN

We benchmarked the performance of RLN vs. purely-data-driven network structures including CARE, RCAN and DDN, which have demonstrated excellent performance in image restoration. The parameters used in training these neural networks are summarized in **Supplementary Table 4**.

The CARE implementation was downloaded from https://github.com/CSBDeep/CSBDeep and networks trained according to their instructions (http://csbdeep.bioimagecomputing.com/doc/). According to the default settings, the number of resolution levels of the U-Net architecture was set to 2, each level in the down-scaling step and the up-scaling step had 2 convolutional layers, the number of convolutional filters for first resolution level was set to 32, and the convolution kernel size was (3 × 3 × 3). The total number of parameters is almost 1 million. During training, the training batch size was set to 4.

For the studies employing RCAN, we used our recently developed 3D RCAN model (https://github.com/AiviaCommunity/3D-RCAN), consisting of 5 residual groups (RG) with each RG containing 5 residual channel attention blocks (RCAB). As default, we used only two convolutional layers in each RCAB. Since the convolution kernel size is (3 × 3 × 3) and the convolution channel number is mostly set as 32, the total number of parameters is over 1 million.

For DenseDeconNet, we used our published single-input neural network (https://github.com/eguomin/regDeconProject/tree/master/DeepLearning) based on three dense blocks. Here we improved the image preprocessing steps by adding online data augmentation and percentile-based normalization. During training, the training batch size was set to 4.

### Training and testing

All networks (RLN, CARE, RCAN and DDN) were implemented with the Tensorflow framework version 1.14.0 and python version 3.6.2 in the Ubuntu 16.04.4 LTS operating system. Training and testing were performed on a computer workstation equipped with 32 GB of memory, an Intel(R) Core(TM) i7 – 8700K, 3.70 GHz CPU, and two Nvidia GeForce GTX 1080 Ti GPUs, each with 24 GB memory. Further details of training and test datasets are summarized in **Supplementary Table 1**.

With this workstation, the maximum size of the input data that RLN can be applied to is 320 MB in 32-bit format. For input data sizes that exceed this limit (e.g., the large cleared-tissue data shown in **Fig. 2d, e, g-i**), our Python-based processing code can automatically crop the volume into several subvolumes, feed them into the RLN network, and stitch the predictions back together. In detail, assuming a data with size W x H x D voxels, we first set the depth d of subvolume as:

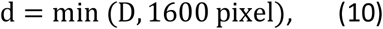

then calculate the width w and height h of subvolume as:

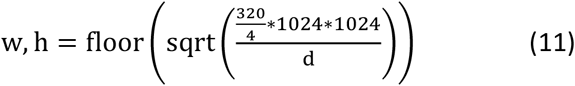

We set the overlapping number voxels in neighboring subvolumes as 24 and use the linear_ramp function (NumPy function) to stitch the overlapped regions. This cropping and stitching procedure is the same as that used in 3D RCAN^11^.

### Quantitative analysis

For all datasets, we selected several volumes (3 - 9) to evaluate the structural similarity index (SSIM) and Peak Signal to Noise Ratio (PSNR) on normalized network outputs and ground truths with MATLAB (Mathworks. R2019b), and then computed the mean value and standard deviation of these volumes. **Supplementary Table 2** summarizes these values.

The Signal to Noise Ratio (SNR) of simulated noisy phantoms (represented as noiseless signal *S* + different levels of noise *N*_*a*_, **Supplementary Fig. 6**) were computed as:

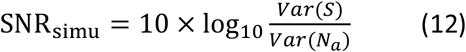

*Var*(·) was used to compute the variance of the volumes. The estimation of SNR of iSIM data (**Supplementary Fig. 16**) is the same as used in our earlier 3D RCAN work^11^:

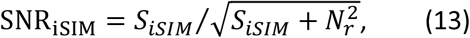

where *S*_*iSIM*_ is the observed, background-corrected signal in photoelectrons (0.46 photoelectrons per digital count) and *N*_*r*_ is the read noise (1.3 electrons according to the manufacturer).

### Sample preparation

For live cell imaging, the U2OS cells were cultured and maintained at 37°C and 5% CO_2_ on a BIO-133 bottomed well plate^20^ for diSPIM imaging (**Fig. 2a-c**), on a #1.5 coverslip (VWR, 48393-241) for diSPIM imaging (**Supplementary Video 1)**, or on glass bottom dishes (Mattek, P35G-1.5-14-C) for iSIM imaging (**Fig. 3, Supplementary Figs. 12, 13, 16**), in 1 mL of DMEM medium (Lonza, 12-604F) containing 10% fetal bovine serum. At 40-60% confluency, cells were transfected with 100 μL of 1X PBS containing 2 μL of X-tremeGENE HP DNA transfection reagent (Sigma,6366244001) and 2 μL plasmid DNA (300-400 ng/μL) and then maintained at 37°C, 5 % CO_2_ for 1-2 days before image acquisition. Cell endoplasmic reticulum (ER) was labeled by ERmoxGFP (Addgene, 68072), mitochondria labeled by mEmerald-Tomm20-C-10 (Addgene, 54281), Golgi apparatus labeled by GalT-GFP (plasmid was a gift from the Patterson Lab, NIH, NIBIB), and lysosomes labeled by Lamp1-EGFP (plasmid a gift from the Taraska Lab, NIH, NHLBI).

For fixed cell imaging **(Fig. 4a-c, Supplementary Fig. 14)**, U2OS cells were cultured on a glass bottom dish and fixed in 4% paraformaldehyde/PBS mixture at room temperature (RT) for 15 minutes, then permeabilized by 0.1% Triton X-100/PBS solution at RT for 2 minutes. Cells were rinsed 3 times by 1X PBS and labeled with 500 μL of 1:100 anti-alpha Tubulin primary antibody (Thermo Fisher Scientific, 322500), 1:200 anti-Tomm20 primary antibody (Abcam, Cat. # 78547), and 1:100 Alexa Fluo 647 Phalloidin (Thermofisher, A22287) in 1X PBS at RT for 1 hour. Labeling mixture was washed away in 1X PBS for 3 times, 1 minute for each time. Cells were then labeled with 500 μL of 1:500 Alexa-488 conjugated donkey anti-rabbit secondary antibody (Invitrogen, Cat. # A21206), 1:500 Alexa-568 conjugated goat anti-mouse secondary antibody (A-11004), and 1 μg/mL DAPI (Thermofisher, D1306) in 1X PBS at RT for 1 hour. After immunolabeling, cells were washed 3 times (1 minute for each time) in 1X PBS.

The mouse brain sample imaged with 0.4/0.4 NA diSPIM (**Fig. 2d**) was prepared using the iDISCO+ procedure and published previously^8^. The brain from an adult mouse with vasopressin receptor 1B Cre X Ai9 was fixed by trans-cardiac perfusion with 4% paraformaldehyde, then dehydrated through a methanol series, rehydrated, immunolabeled with an antibody for tdTomato (Rabbit anti-RFP, Rockland Antibodies and Assays, Cat. # 600-401-379) and an Alexa 555 secondary antibody (Invitrogen, Cat. # A27039). Before imaging with cleared tissue diSPIM^8^, the tissue slab was dehydrated with a methanol series, and dichloromethane before equilibration in dibenzyl ether (Sigma, Cat. # 108014).

For the cleared mouse brain samples (**Fig. 2g, Supplementary Fig. 8b, c, Supplementary Video 2)**, fixed adult mouse brain expressing tdTomato in axonal projections from the area of a stereotaxic injection of AAV was cleared using SDS and equilibrated in CUBIC-R^32^. 2 mm thick coronal slabs were sectioned and held in a sample chamber custom designed for the CT-DISPIM.

Nematode strains included BV24 ([*ltIs44* [*pie-1*p-mCherry::PH(PLC1delta1) + *unc-119*(+)]; *zuIs178* [(*his*-72 1 kb::HIS-72::GFP); *unc-119*(+)] V], **Supplementary Figs. 10a-b**,**11b, Supplementary Video 3**), od58 (*ltIs38 [pie-1p::GFP::PH(PLC1delta1) + unc-119(+)]*, **Fig. 1b, Supplementary Figs. 10c-f, 11a**), and DCR6268 ([*pttx-3b*::SL2::Pleckstrin homology domain::GFP::unc-54 3’UTR + *pelt-7*::mCh::NLS::*unc-54* 3’UTR]), **Fig. 4d-f, Supplementary Fig. 15**). All worms were cultivated at 20°C on nematode growth medium plates seeded with a lawn of *Escherichia coli* strain OP50. Embryos were dissected from gravid adults, placed on poly-L-lysine-coated coverslips and imaged in M9 buffer, as previously described^33^.

### Simulation of phantom objects

To evaluate the quality and performance of our network, we generated 3D phantom objects consisting of three types of structures in MATLAB (Mathworks, R2019b, with the Imaging Processing Toolbox) for ground truth: dots, solid spheres, and ellipsoidal surfaces (**Supplementary Fig. 2**). Each phantom was composed of 100 solid spheres, 100 ellipsoidal surface and 400 dots, randomly located in a 128 × 128 × 128 volume. The 100 solid spheres were generated with random intensity (50-850 counts) and random diameter (4-8 voxels); The 100 ellipsoidal surfaces were generated with random intensity (50-850 counts), random diameter along different axes (4-8 voxels) and random thickness (1-2 voxels); the 400 dots were generated with random intensity (50-850 counts) and random extent along each direction (1-3 voxels). The background value was set to a constant at 30 counts.

Noiseless input volumes were generated by convolving the ground truth data with different PSFs (**Supplementary Fig. 2**). Four types of PSFs were used, including: the system PSF for the 0.8/0.8 NA diSPIM which has 3-fold larger axial extent compared to its lateral extent^19^ for the generalization test on embryo nuclei and membrane data (**Supplementary Fig. 11**); the system PSF of iSIM^11^ for the generalization test of ER volumes (**Fig. 3e, Supplementary Fig. 13d-f**), the system PSFs of the widefield microscope (Olympus, UPLXAPO60XO, 60X, NA = 1.42 oil objective) for the generation test of the four color fixed U2OS cells (**Fig. 4a-c**), and the system PSF of the widefield microscope (Olympus UPLSAPO60XWPSF, 100X, NA = 1.35 silicon oil lens) for the generation test of *C. elegans* embryo expressing ttx-3B-GFP (**Fig. 4d-f**). Noisy images were then obtained by adding different levels of Gaussian and Poisson noise.

The 3D human brain phantom was downloaded from the Zubal Phantom website^34^ (http://noodle.med.yale.edu/zubal/data.htm, **Fig. 1e-f, Supplementary Fig. 4**). The simulated spherical phantoms ground truths were generated with ImgLib2^18^ and blurred with a 3D Gaussian kernel with standard deviation set to 2, the maximum radius of the spheres was set at seven pixels and the intensity range to 80–255 (**Fig. 1d, Supplementary Figs. 5, 7**). Network inputs of these structures (**Fig. 1d, e**) and their corresponding training data were blurred with the system PSF of the 0.8/0.8 NA diSPIM.

### DiSPIM data acquisition and processing

A fiber-coupled diSPIM^16^ with two 40x, 0.8 NA water objectives (Nikon Cat. # MRD07420), resulting in a pixel size of 162.5 nm, was used to image the U2OS cell transfected with mEmerald-Tomm20-C-10 (**Fig. 2a, Supplementary Video 1**), transgenic embryos strain od58 expressing GFP-membrane (**Fig. 1b, Supplementary Figs. 10c-f, 11a**) and BV24 expressing GFP-nuclei (**Supplementary Figs. 10a-b, 11b, Supplementary Video 3**). For cellular imaging, 50-200 dual-view volumes (60 planes, 1 μm inter-plane spacing in each view) were acquired with 3 s intervals; for embryo imaging, dual-view stacks (50 planes at 1 μm spacing per view) were acquired at 1-minute intervals for 291 minutes. Dual-view data were registered and jointly deconvolved with ImageJ plugin diSPIM Fusion^8^ for the ground truth, with 10 iterations.

### DiSPIM cleared tissue acquisition and processing

Cleared tissue image data in **Fig. 2d** was acquired on a fiber-coupled diSPIM that was modified for cleared-tissue imaging by incorporating elements of the commercially available ASI DISPIM and DISPIM for Cleared Tissue (CT-DISPIM)^8^. We used a pair of Special Optics 0.4-NA multi-immersion objectives (ASI, 54-10-12). The cleared mouse brain volumes were acquired by moving the stage (2 μm step size, total 4800 frames with 2048 × 2048 pixels) in a raster pattern with the aid of the ASI diSPIM Micromanager plugin (http://dispim.org/software/micro-manager). Image data for **Fig. 2g** were acquired on a dedicated, commercial ASI CT-DISPIM equipped with a pair of Special Optics 0.7-NA multi-immersion objectives (ASI, 54-12-8). Using the DISPIM plugin in Micromanager, we set up a multi-position acquisition in light sheet mode with unidirectional stage scan. Image FOV was set to 1536 × 1536 pixels to avoid geometric distortions near the edge of the full FOV (2048 × 2048). Five y positions and 2 z positions were acquired with 15% overlap, each position was a stack of 1573 images with a stage step of 1.414 μm.

Dual-view data were registered and jointly deconvolved based on Wiener-Butterworth filter back projector (1 iteration) for the ground truth, using MATLAB (Mathworks, R2019b, with the Imaging Processing and Parallel Computation Toolboxes)^8^ on a computer workstation equipped with Intel(R) Xeon(R) W-2145 CPU @ 3.70GHz and Nvidia Quadro P6000 with 24 GB memory.

For joint deconvolution of cleared mouse brain samples in **Fig. 2g**, running on National Institutes of Health (NIH) Biowulf cluster, we modified the code to meet the HPC requirements for job scheduling. The 28-core “gpu” queue for Biowulf (28 × 2.4 GHz Intel E5-2680v4 processor, 4 x NVIDIA P100 GPUs, 16 GB VRAM, 3584 cores) was used for computing.

### iSIM data acquisition and processing

A home built iSIM system^23^ with 60X, NA = 1.2 water objective (Olympus UPLSAPO60XWPSF) and an sCMOS camera (PCO, Edge 5.5), resulting in a pixel size of 55 nm, was used to image the U2OS cells (**Fig. 3, Supplementary Figs. 12, 13, 16**). All raw volumes were background subtracted and deconvolved using the Richardson-Lucy algorithm with 15 iterations as the ground truth.

### Wide field data acquisition and processing

Wide-field fixed U2OS images (**Fig. 4a-c, Supplementary Fig. 14**) were acquired by a home built wide field microscope with a 60X, NA=1.42 oil objective (Olympus, UPLXAPO60XO) and a pixel size of 266 nm. Widefield *C*. elegans embryos (**Fig. 4d-f, Supplementary Fig. 15**) were acquired with a 100X, NA = 1.35 silicon oil lens (Olympus UPLSAPO60XWPSF) and a pixel size of 111 nm.

## Notes

### Competing Interest Statement

The authors have declared no competing interest.

