## Supplemental Materials for "Incorporating the image formation process into deep learning improves network performance in deconvolution applications"

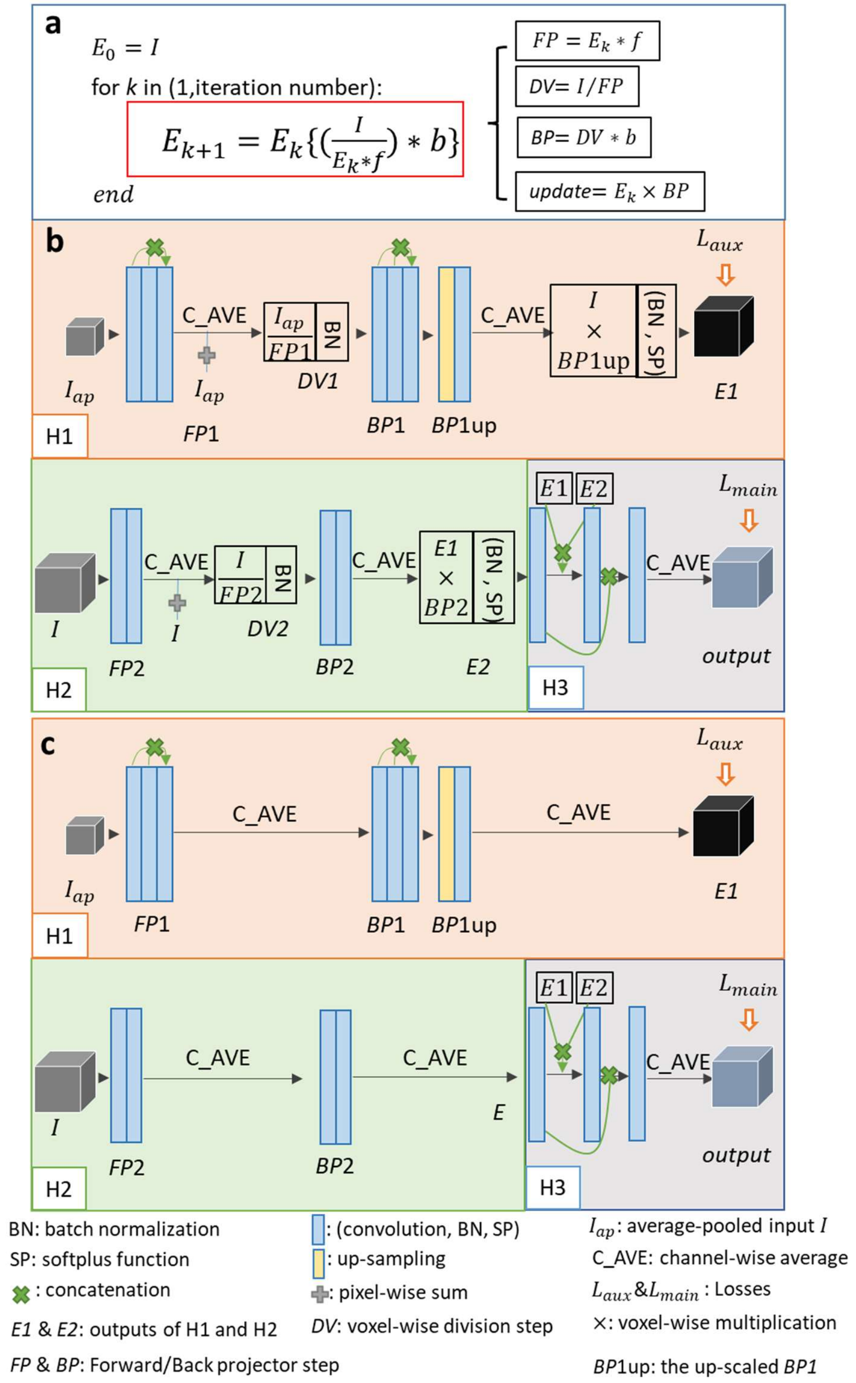


**Supplementary Fig. 1, Decomposition of RL deconvolution iteration and the internal structure of RLN and RLN-a. a)** RL deconvolution can be decomposed into four parts: forward projector (*FP*) function, division (*DV*) step, back projector (*BP*) function and *update* step. **b)** Schematic of RLN consisting of three parts: down-scale estimation starting from the average-pooled input image, H1; original-scale estimation starting with original-scale input image, H2; and merging/fine-tuning, H3. H1 and H2 are inspired by the RL deconvolution update formula, which mimic the unmatched forward/back project steps. **c)** Schematic of RLN-a, without the *DV* and *update* steps in RLN. See **Methods** for more detail.

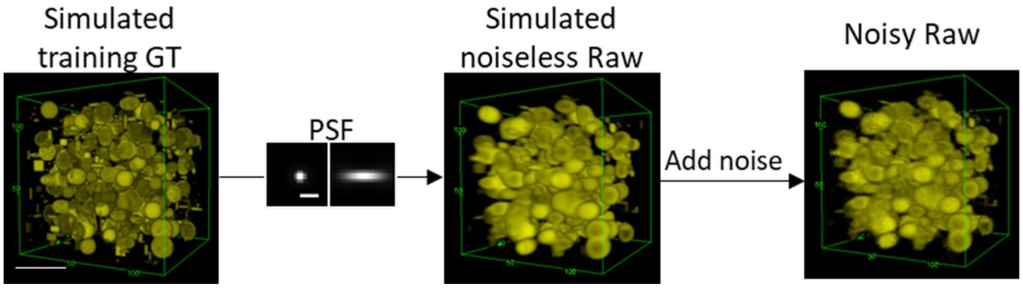


**Supplementary Fig. 2, The training simulated data generation process.** Simulated training ground truth (GT) consists of dots, solid spheres, and ellipsoidal surfaces. The noiseless raw data is generated by convolving the ground truth data with the PSF. Scale bar: 5 µm.

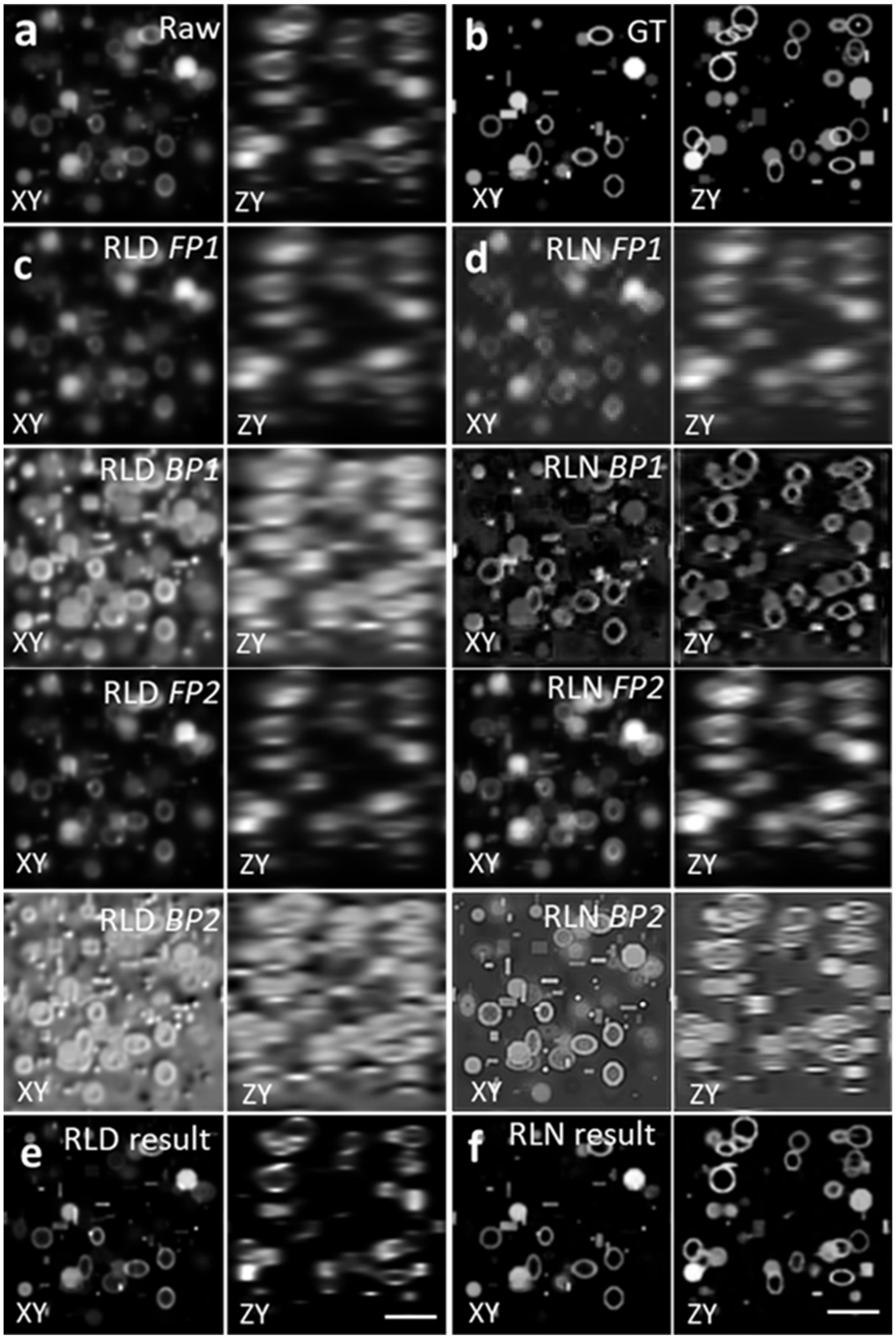


**Supplementary Fig. 3**, **Comparing RLD and RLN on a phantom object consisting of dots, solid spheres, and ellipsoidal surfaces**. Lateral (XY) and axial (ZY) views are presented in each case. **a)** Raw input, i.e., blurry image. **b)** Ground truth object. **c)** Intermediate output of RLD. *FP1* and *BP1* are the forward projector function and backward projector function at iteration 1, and *FP2, BP2* at iteration 20. **d)** Intermediate steps of RLN. *FP1, BP1, FP2, BP2* are the steps in H1 and H2 shown in the neural network (**Supplementary Fig. 1**). The similarities between the RLD and RLN intermediate steps indicates the RLN internal feature maps are interpretable. **e)** Final RLD result after 40 iterations. **f)** RLN result, which is much closer to the ground truth (SSIM 0.95, PSNR 31.1) than RLD (SSIM 0.67, PSNR 19.4). Scale bars: 5μm.

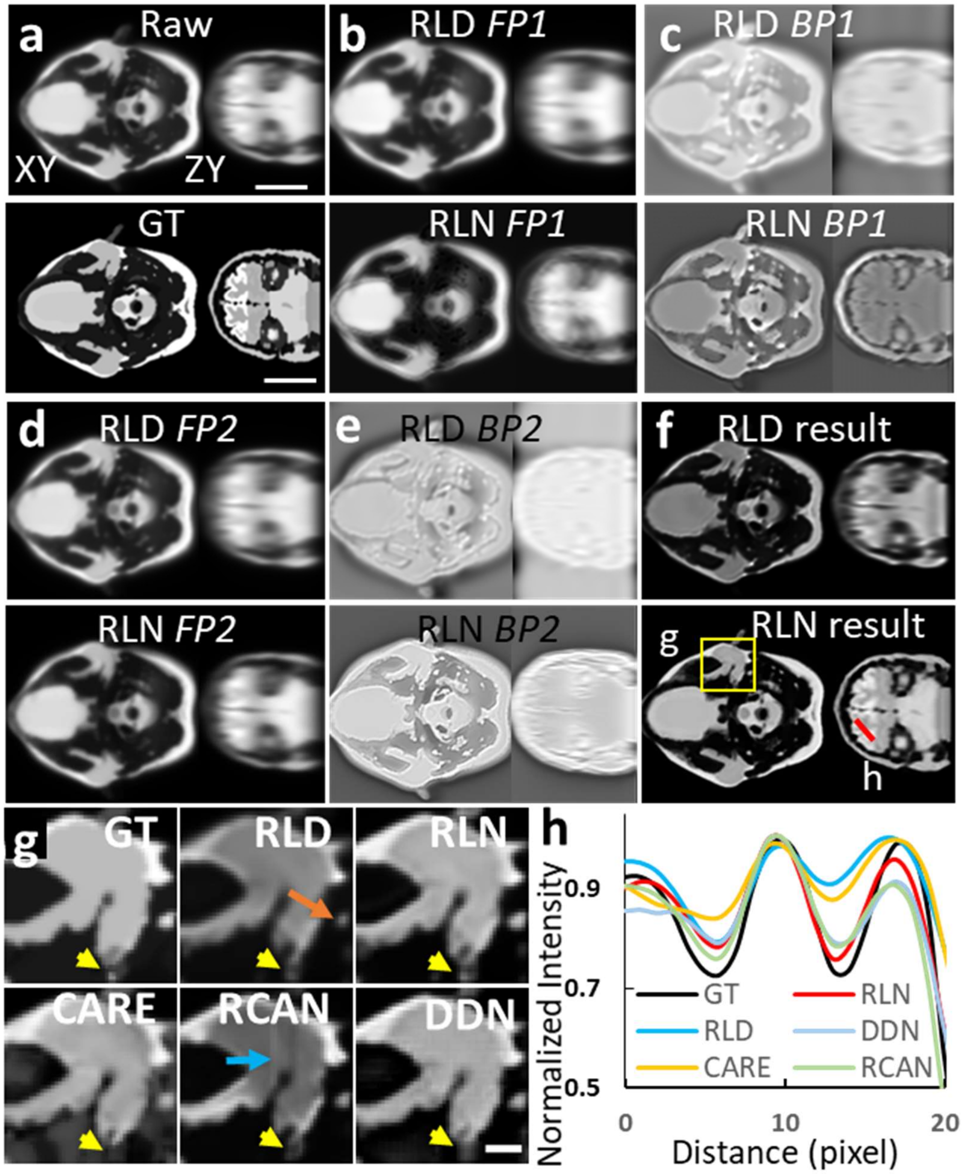


**Supplementary Fig. 4**, **Interpretability and generalization of RLN as assessed on human brain phantom. a)** Blurry raw input (top row) and ground truth (GT, bottom row). **b-e)** Intermediate output of RLD (top row) and RLN (bottom row) at *FP1* **(b)**, *BP1* **(c)**, *FP2* **(d)**, and *BP2* **(e)**. Similarities in output between RLN and RLD indicates the interpretability of the RLN. **f)** Results of RLD (top row) and RLN (bottom row). The RLN result is noticeably closer to the ground truth (SSIM 0.89, PSNR 24.4) than RLD (SSIM 0.72, PSNR 16.9). Lateral (left) and axial (right) views are shown in **a-f)**. **g)** Magnified view of the yellow rectangle in **f)**, comparing the ground truth, RLD and the predictions from RLN, CARE, RCAN and DDN, with arrows highlighting features showing RLN provides better restoration than the other methods, see also **Fig. 1g** for quantification. **h)** Line profile along the red line in **f)**, showing RLN is closer to the ground truth than RLD and other networks. All models were trained with the phantom objects consisting of dots, solid spheres, and ellipsoidal surfaces. See also **Fig. 1f**. Scale bars: **a-f)** 50 pixels**, g)** 10 pixels.

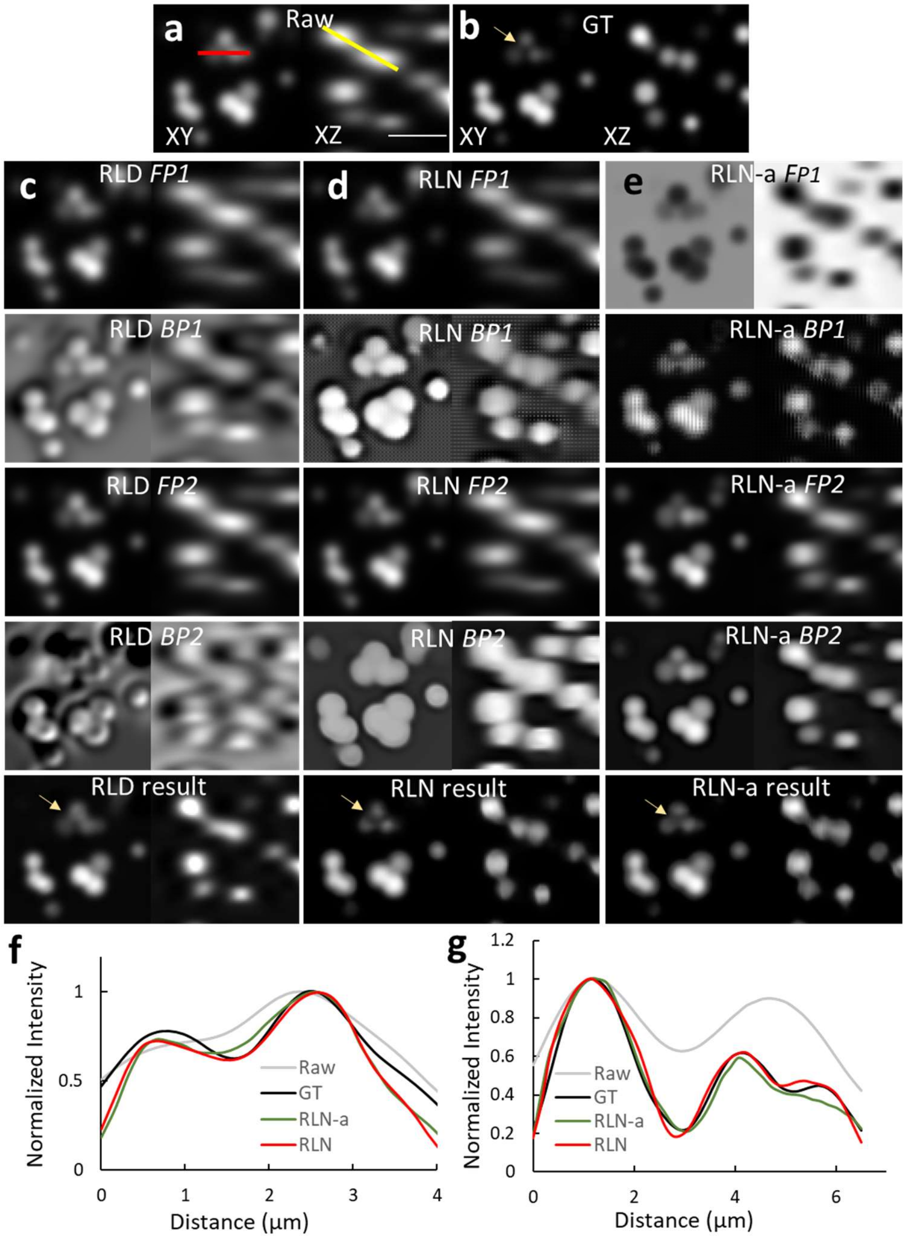


**Supplementary Fig. 5, Comparing RLD, RLN, and RLN-a performance on simulated beads, using training data from synthetic mixed structures**. Lateral (left) and axial (right) views are shown in each case. **a)** Raw input. **b)** Ground truth. **c-e)** Intermediate output of **c)** RLD, **d)** RLN, and **e)** RLN-a. RLN and RLN-a models were trained with the phantom objects consisting of dots, solid spheres, and ellipsoidal surfaces. Dim details are better restored with RLN than RLD and RLD-a (yellow arrows). RLN shows better recovery (SSIM 0.97, PSNR 35.7) than RLN-a (SSIM 0.94, PSNR 34.0). Also, the intermediate output of RLN, appear visually closer to the intermediate steps of RLD, particularly the *FP1* step. These results suggest that the additional network structure in RLN (i.e., the *DV* and *update* steps) aids in network generalizability. **f, g)** Line profiles along the red and yellow lines in the XY and ZY views in **a),** showing RLN is closer to the ground truth than RLD and RLN-a. Scale bar: 5 µm.

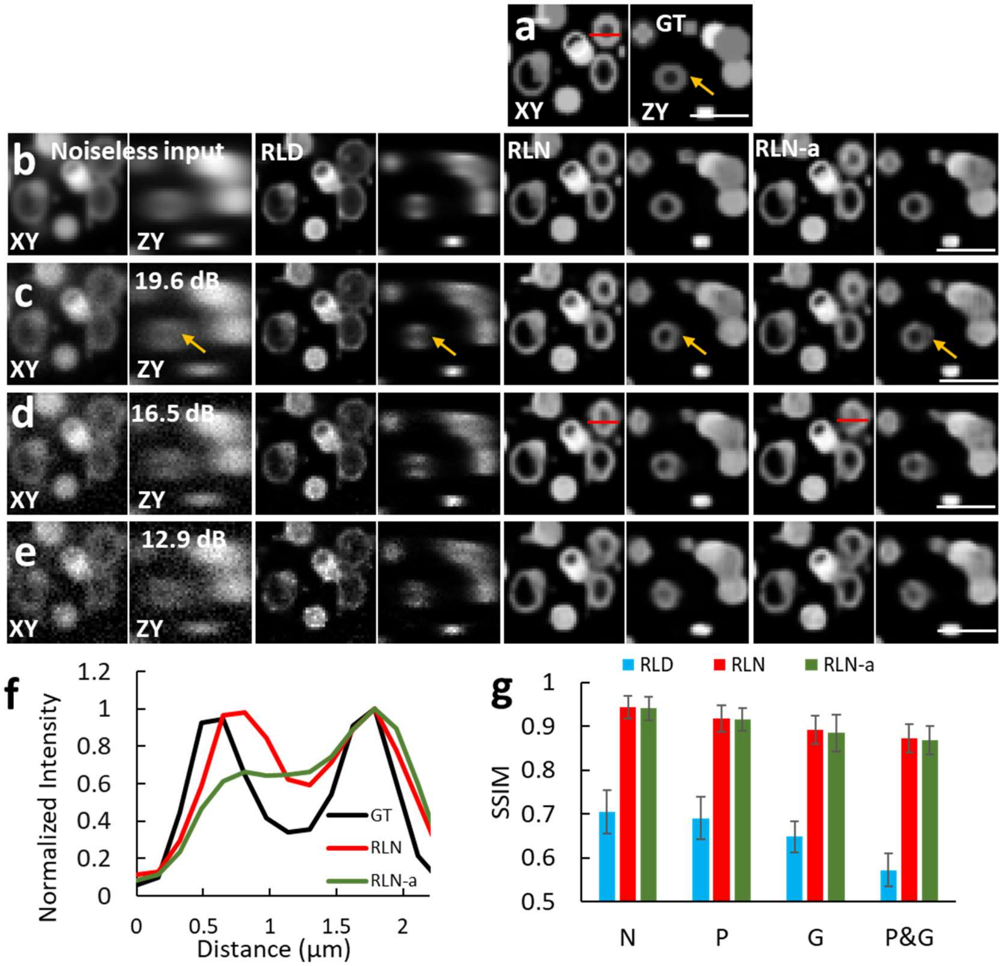


**Supplementary Fig. 6, Deconvolution quality as a function of input SNR, comparing RLD, RLN and RLN-a trained with synthetic mixed structures dataset with corresponding SNR. a)** Ground truth. **b-e)** Left to right columns show output of RLD, RLN, and RLN-a in lateral (left) and axial (right) views as a function of **b)** no noise (N); **c)** Poisson noise (P); **d)** Gaussian noise (G); **e**) mixed Poisson and Gaussian noise (P&G). **f)** Line profiles of the red lines in **a)**, **d)** showing that RLN prediction is closer to ground truth than RLN-a. **g)** SSIM and PSNR values of RLD, RLN, and RLN-a as a function of different noise types. In all cases, SSIM and PSNR decrease as noise increases, and RLN and RLN-a show considerably higher SSIM and PSNR than RLD at all noise levels. Means and standard deviations from N=9 volumes are shown, yellow arrows highlight example structure best resolved in RLN compared to RLN-a and RLD. Scale bars: 3 µm.

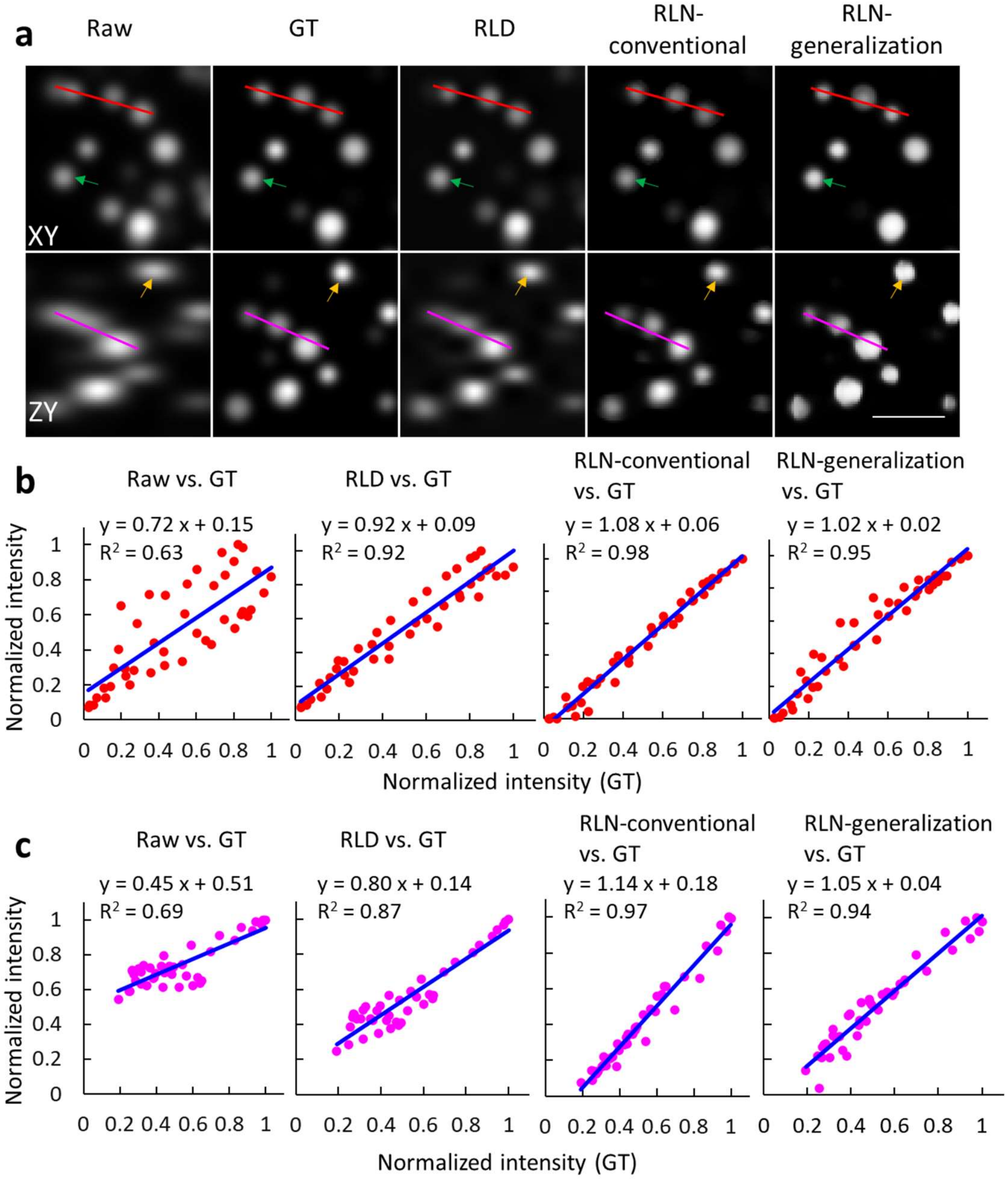


**Supplementary Fig. 7, RLN performance on simulated bead samples, comparing RLD, conventional testing, and generalization. a)** Raw image, ground truth (GT), RLD result, and RLN predictions in lateral (top) and axial (bottom) views. The model for generalization was trained with phantom objects consisting of mixed dots, solid spheres, and ellipsoidal surfaces, whereas the model for the conventional test used the same type of training data as the test data (simulated beads). Although RLN always outperforms RLD, the generalization result slightly distorts and sharpens bead shapes compared to the ground truth and conventional testing result (green and yellow arrows). Scale bar: 5 µm. **b)** Normalized intensity of raw, RLD, and RLN predictions (y axis) vs. normalized intensity of ground truth (x axis) taken along the red line shown in the lateral view in **a)**. The scattered red dots indicate pixel intensities, the solid blue line is the linear fit to the data, and the insets display the fitting equation and the square of the correlation coefficient ($R^2$). **c)** as in **b)**, but for the magenta line shown in the axial view in **a)**. These data suggest that the RLN linearity is better than RLD.

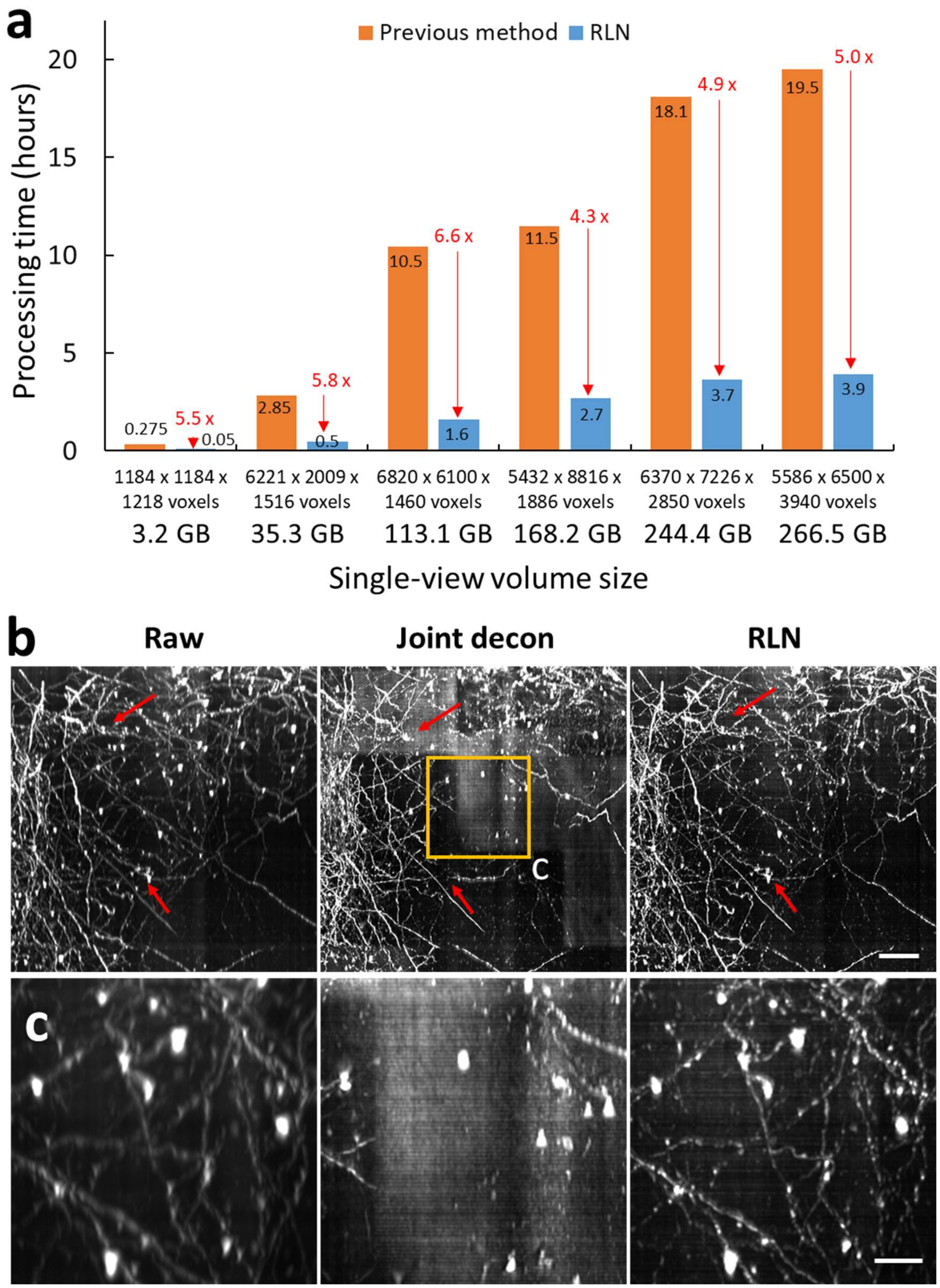


**Supplementary Fig. 8**, **RLN outperforms the previous processing pipeline for reconstructions of large, cleared tissue datasets imaged with diSPIM. a)** Processing time of RLN vs. previous processing pipeline (i.e., cropping, registration, joint deconvolution, and stitching) on differently sized volumes. Data are 16-bit. The 3.2 GB and 168.2 GB data were from the cleared brain tissue shown in **Fig. 2**, others were from the previously published cleared tissue samples and reprocessed with RLN. As shown, RLN accounts for a 4- to 6-fold speed improvement. **b)** Comparisons of single-view raw, dual-view joint deconvolution and RLN predictions from the subvolume shown in **Fig. 2g.** Red arrows indicate artifacts in dual-view joint deconvolution result, relative to the raw input and RLN prediction. We suspect these artifacts are likely due to failures of registration between the two raw views. **c)** Magnified view of rectangular regions in **b)**, further demonstrating the poor joint deconvolution results. Scale bars: **b)** 50 µm; **c**) 20 µm.

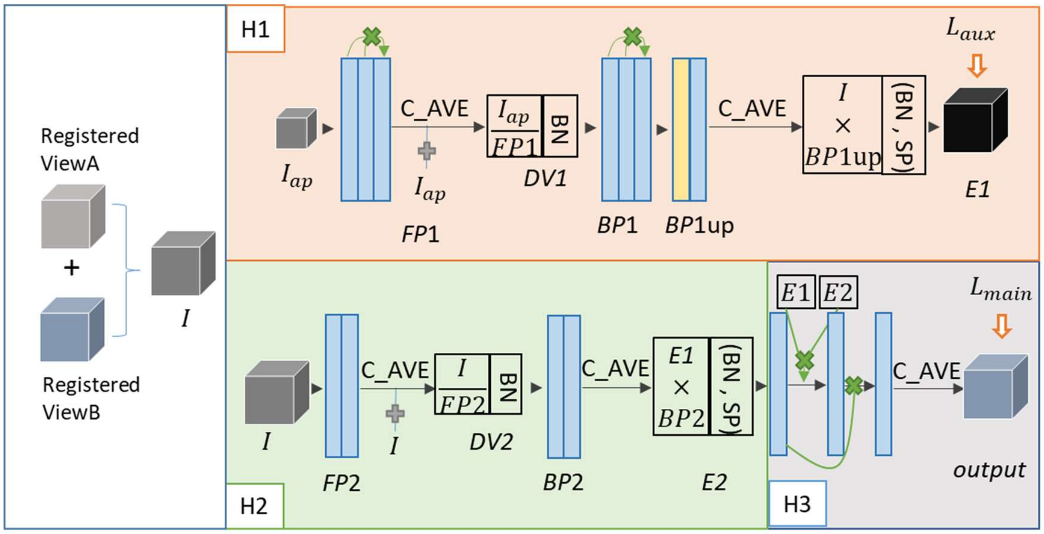


**Supplementary Fig. 9, Dual-input RLN.** The registered dual views are merged by averaging before applying the network.

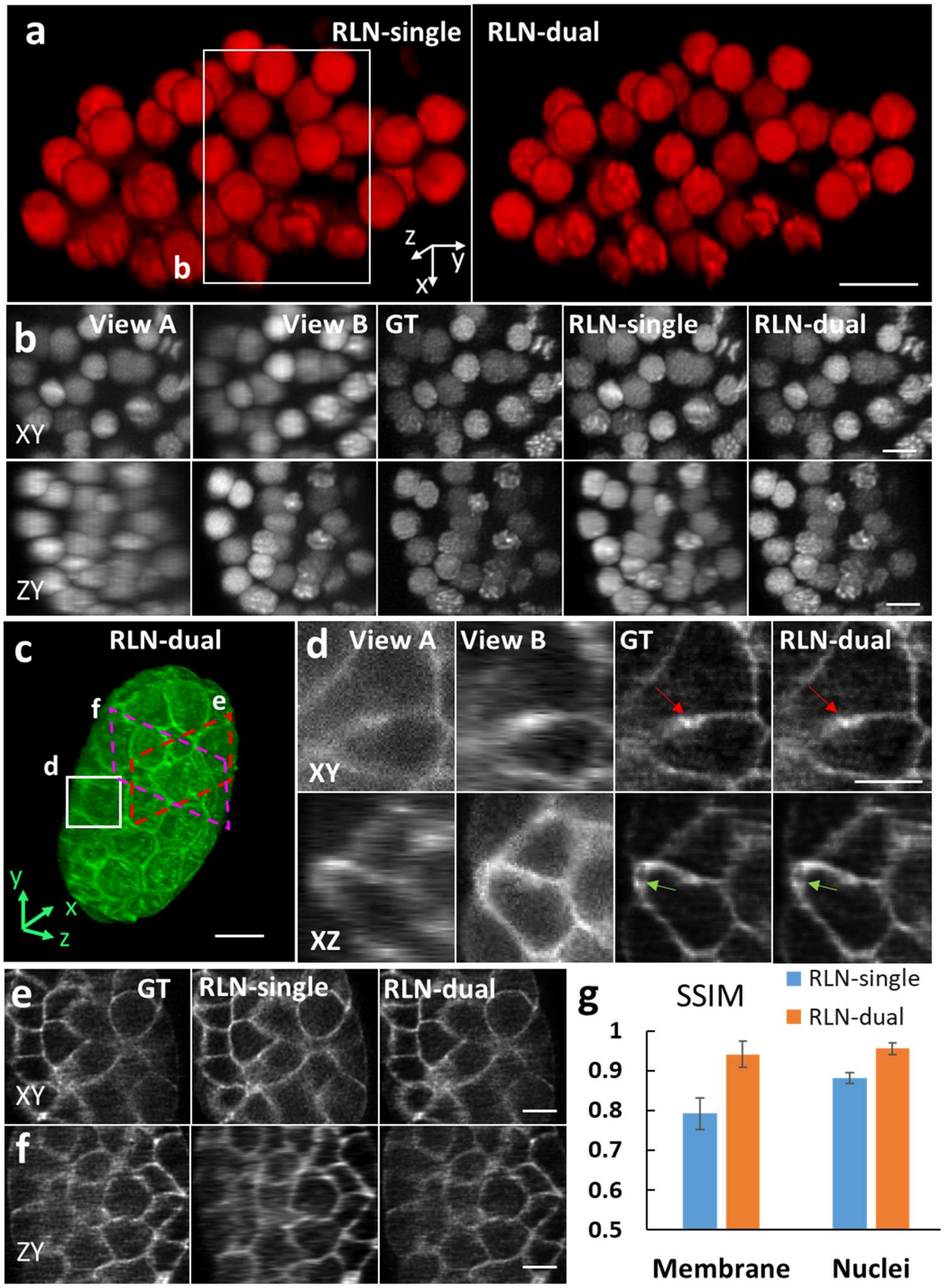


**Supplementary Fig. 10, Dual-input RLN improves axial resolution relative to single-input RLN on scattering samples. a)** 3D rendering of nuclei expressed in live *C. elegans* embryo collected by diSPIM and processed with single-input RLN and dual-input RLN. **b)** Higher magnification of white rectangle in **a),** comparing raw view A, raw view B, joint deconvolution of views A/B, single-input RLN and dual-input RLN predictions. The dual-input RLN uses the information available in both views to achieve resolution isotropy, providing near-identical output to the ground truth. **c)** 3D rendering of membrane marker expressed in live *C. elegans* embryo, raw data collected by diSPIM and processed with dual-input RLN. **d)** Higher magnification of white rectangle in **c),** comparing raw view A, raw view B, joint deconvolution of views A/B, and dual-view RLN predictions. Dual-input RLN uses complementary dual-view information to enable near-isotropic resolution, providing results near-identical to the ground truth (red and green arrows). **e, f)** Higher magnification of dashed red and magenta parallelograms in **c),** comparing dual-view deconvolution ground truth, single- and dual-view RLN predictions. The dual-input RLN better recovers axial resolution than single-input RLN. **g)** SSIM analysis for data displayed in **b)** and **e),** showing that RLN-dual better recovers the signals than RLN-single when the views are contaminated by scattering. Means and standard deviations are obtained from N = 9 volumes for membranes and nuclei. Scale bars: **a, b, d, e, f)** 5 µm, **c)** 10 µm.

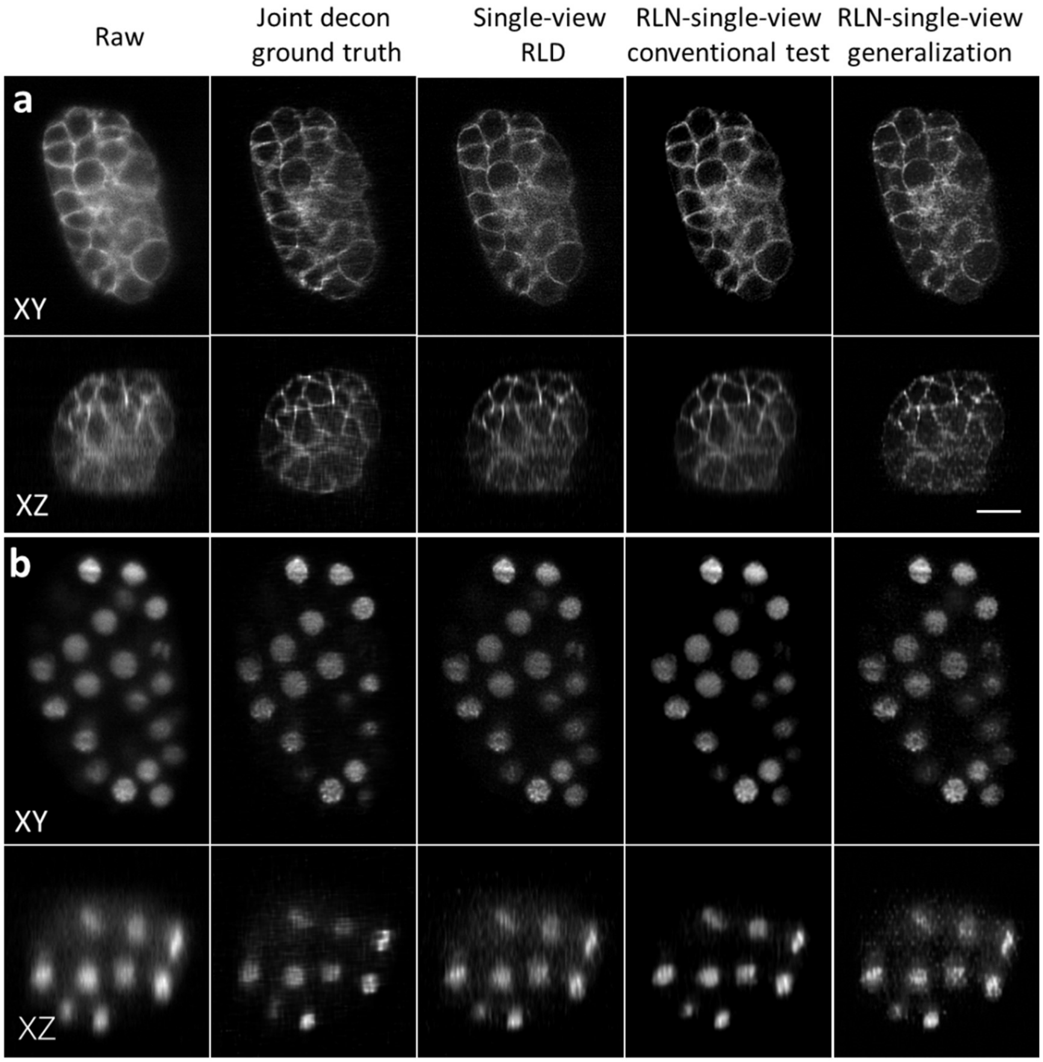


**Supplementary Fig. 11, RLN models trained on synthetic mixed data generalize well to images of *C. elegans* embryos acquired with diSPIM. a)** Membrane and **b)** nuclei results, comparing raw input, joint deconvolution ground truth, single-view RLD, and predictions from RLN with single-view input under conventional testing (trained on similar images to test data) vs. generalization (models trained with mixed phantoms of dots, solid spheres, and ellipsoidal surfaces). Outputs of single-view RLD, conventional testing, and generalization show close visual resemblance to each other. Quantitative assessments show that the conventional testing results (SSIM-membrane: 0.80, SSIM-nuclei: 0.85, PSNR-membrane: 28.9, PSNR-nuclei: 27.0) is closest to the dual-view joint deconvolution ground truth, while the generalization results (SSIM-membrane: 0.75, SSIM-nuclei: 0.76, PSNR-membrane: 27.3; PSNR-nuclei: 26.1) compare favorably against single-view RLD (SSIM-membrane: 0.74, SSIM-nuclei: 0.73, PSNR-membrane: 27.1, PSNR-nuclei: 25.9). Scale bars: 10 μm.

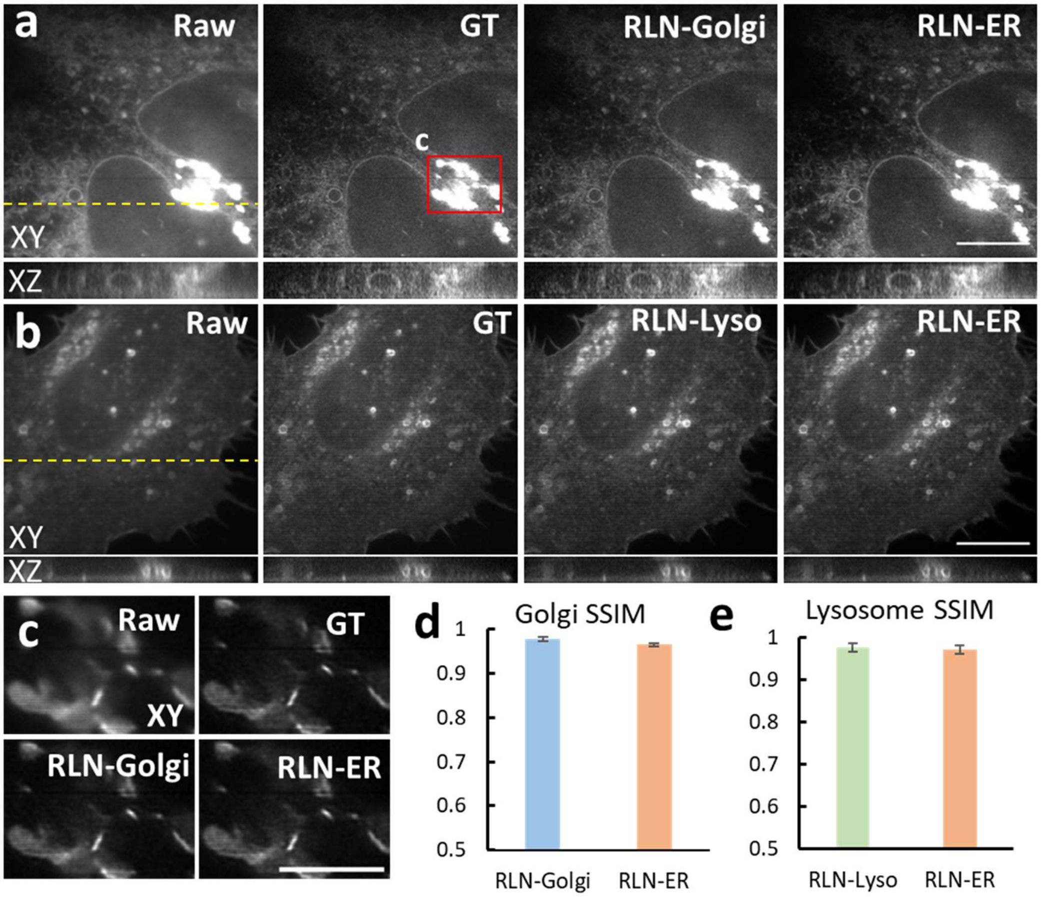


**Supplementary Fig. 12, Generalization ability of RLN tested with ER, Golgi, and Lysosome markers, imaged with iSIM. a)** Lateral and axial views of live U2OS cells expressing GalT-GFP, acquired with iSIM, comparing the raw input, deconvolved ground truth, predictions from Golgi-trained RLN (RLN-Golgi) and ER-trained RLN (RLN-ER). Yellow dashed line indicates the Y position of the XZ plane. **b)** Lateral and axial views of live U2OS cells expressing Lamp1-EGFP, acquired with iSIM, comparing the raw input, ground truth, predictions from lysosome-trained RLN (RLN-Lyso) and ER-trained RLN (RLN-ER). Axial view is taken along yellow dashed line. **c)** contrast-adjusted higher magnification view of red rectangular region in **a)**. **d)** Quantitative SSIM analysis for data shown in **a)**, means and standard deviations are obtained from N = 6 volumes. **e)** Quantitative SSIM analysis for data shown in **b)**, means and standard deviations are obtained from N = 6 volumes. Scale bars: **a, b)** 10 µm, **c)** 5 µm.

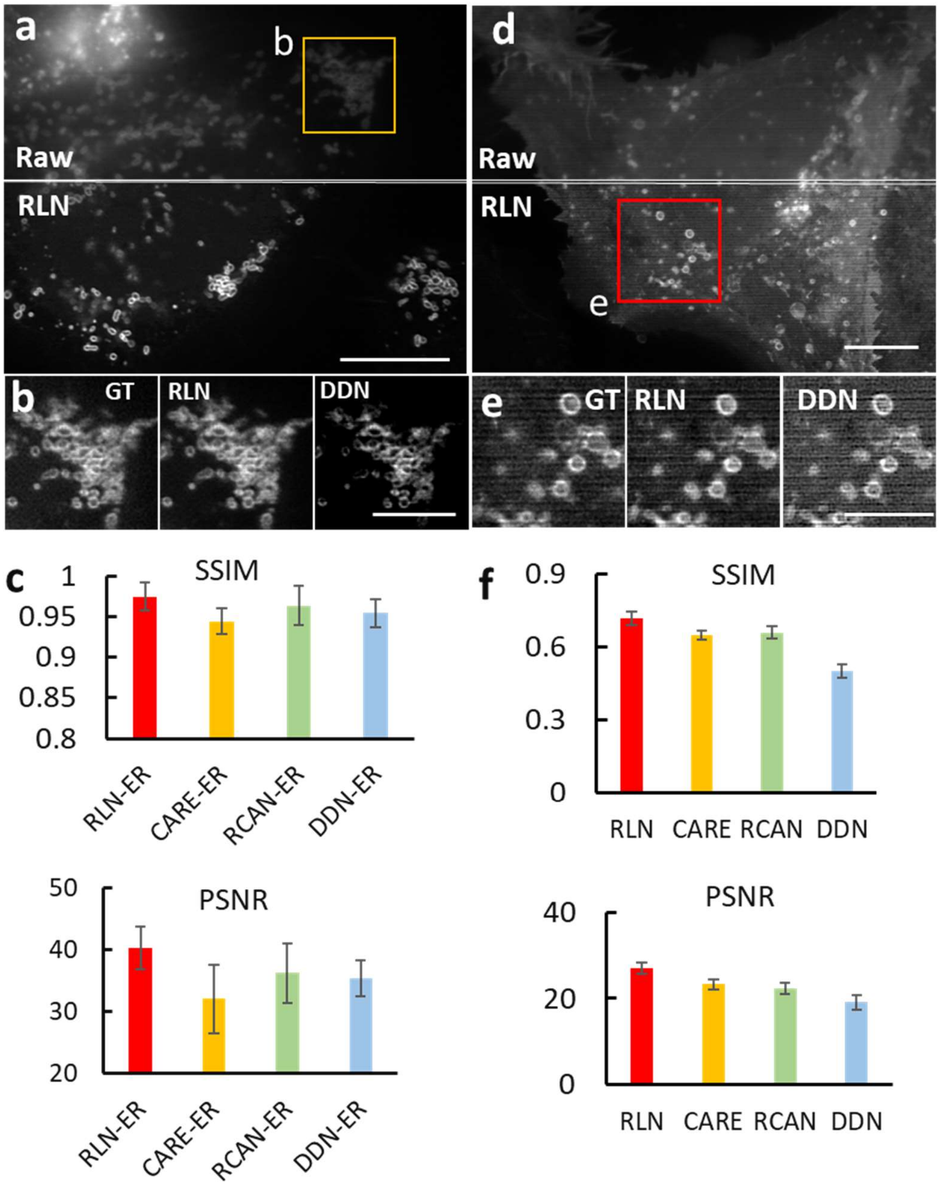


**Supplementary Fig. 13**, **RLN provides better generalization on super-resolution data than other networks. a)** Super-resolved images of live U2OS cells expressing mEmerald-Tomm20-C-10, acquired with iSIM. Top: raw input; bottom: RLN output. **b)** Higher magnification view of yellow rectangular region in **a)**, comparing ground truth, RLN output, and DDN output. The models were trained with ER datasets. **c)** SSIM and PSNR measurements for RLN, CARE, RCAN and DDN for data shown in **a)**, means and standard deviations are obtained from N = 6 volumes. **d)** Super-resolved images of live U2OS cells expressing Lamp1-EGFP, acquired with iSIM. Top: raw input; bottom: RLN output. **e)** Higher magnification of rectangular regions in **d)**, comparing ground truth, RLN output, and DDN output. Models were trained with phantom objects consisting of dots, solid spheres, and ellipsoidal surfaces. **f)** SSIM and PSNR measurements, comparing RLN, CARE, RCAN and DDN for data shown in **d)**, means and standard deviations are obtained from N = 6 volumes. Scale bars: **a, d)** 5 µm, **b, e)** 2 µm.

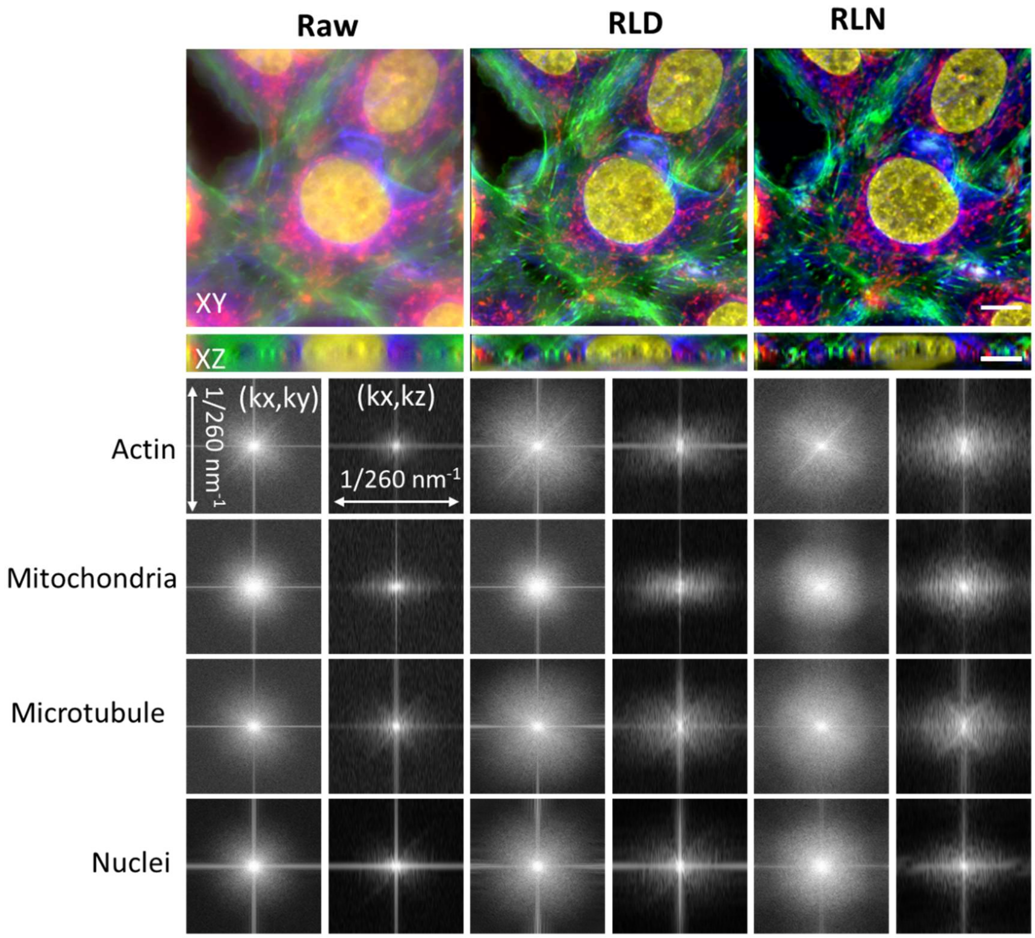


**Supplementary Fig. 14**, **Four color lateral and axial maximum intensity projections and Fourier spectra of a fixed U2OS cell.** Images were acquired by widefield microscopy; here raw input, RLD, and RLN predictions based on a model trained on the synthetic mixed structures are compared. See also **Fig. 4a-c**. Red: mitochondria immunolabeled with anti-Tomm20 primary antibody and donkey α-rabbit-Alexa-488 secondary; green: actin stained with phalloidin-Alexa Fluor 647; Blue: tubulin immunolabeled with mouse-α-Tubulin primary and goat α-mouse-Alexa-568 secondary; yellow: nuclei stained with DAPI. Images and Fourier spectra in axial and lateral views indicate that RLD better recovers resolution than RLD. Scale bars: 20 µm.

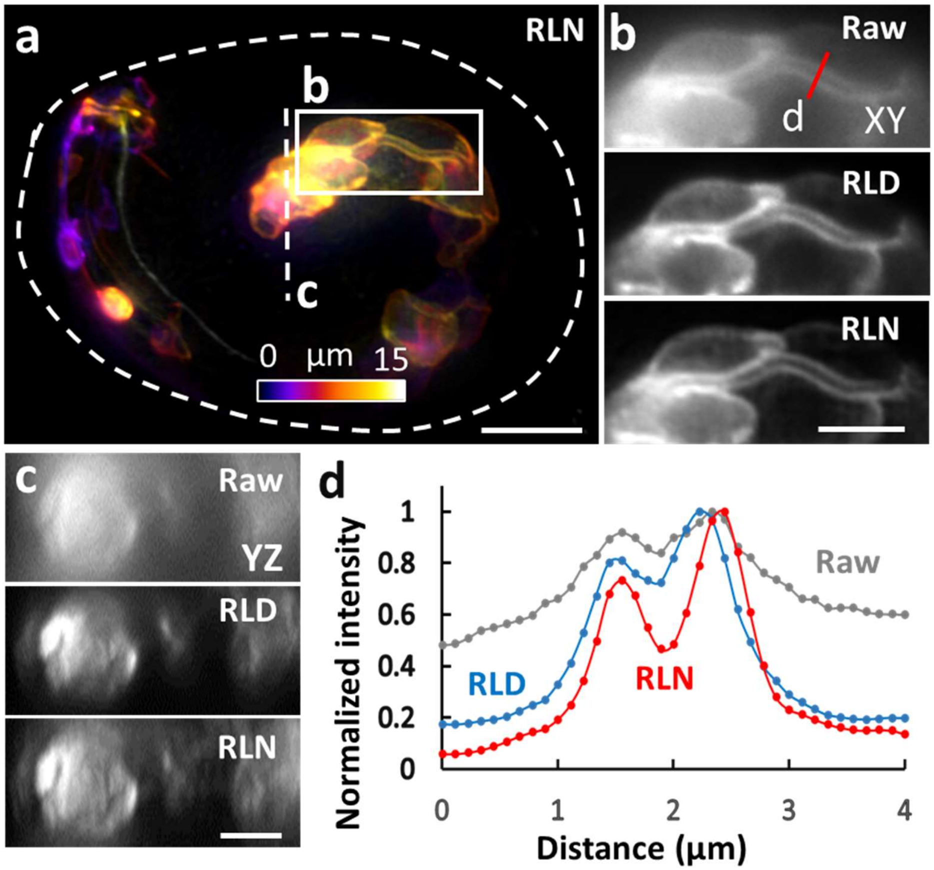


**Supplementary Fig. 15**, **RLN trained with synthetic mixed structures outperforms direct RLD on a *C. elegans* embryo stack. a)** Depth-coded reconstruction of *C. elegans* embryo expressing ttx-3B-GFP, acquired by widefield microscopy, and predicted by RLN based on a model trained on synthetic mixed structures. Dashed line indicates the embryo shape. See also **Fig. 4d-f**. **b, c)** Higher magnification of white rectangle and dashed line in **a)**, comparing the raw input, RLD, and RLN prediction, highlighting the membranes of individual gut cells in lateral **b)** and axial views **c)**. **d)** Line profiles of the red line shown in **b)**, indicating that RLN better resolves the membrane structure than RLD. Scale bars: **a)** 10 µm, **b, c)** 5 µm.

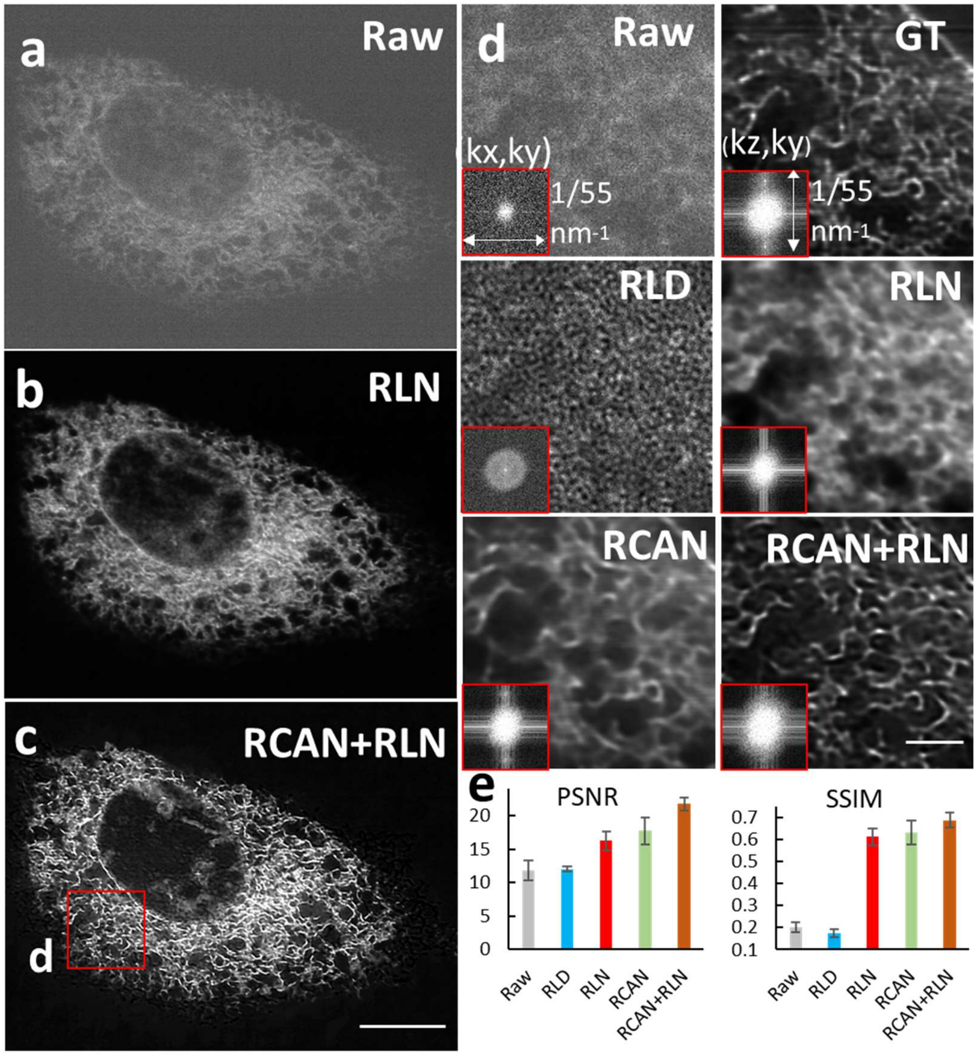


**Supplementary Fig. 16, Comparison of RLD and RLN on super-resolved, low-SNR data from a live U2OS cell expressing ERmoxGFP, acquired with iSIM. a)** Low SNR input, XY view. **b)** RLN output with one step training, i.e., applying a model trained with low SNR raw input and high SNR deconvolved ground truth. **c)** RLN output with two-step deep learning, by first applying a denoising RCAN model, then applying an RLN model to deconvolve the output from the first step. The RCAN model was based on pairs of low/high SNR raw data, the RLN model was based on pairs of high SNR raw data and high SNR deconvolution data. **d)** Higher magnification of red rectangle in **c),** comparing raw input, high SNR deconvolved ground truth, RLD on low SNR raw input, one-step RLN, one-step RCAN (same training data as with one-step RLN, i.e., input is the low SNR raw input and the high SNR deconvolved result is the ground truth), and two-step deep learning with RCAN for denoising and RLN for deconvolution. Insets show Fourier transforms of the data. **e)** Quantitative analysis with PSNR and SSIM for the raw input, RLD, one-step RLN, one-step RCAN, and two-step RCAN+RLN result, means and standard deviations are obtained from N = 6 volumes. Both one-step RCAN and one-step RLN outperform RLD, and the two-step methods further boosts resolution and contrast, indicated by the Fourier spectra shown in the inserts. Scale bars: **a-c)** 10 µm, **d)** 2 µm.

**Supplementary table 1, Summary of RLN training and testing datasets.**

| | Figures | Training dataset | Training data size (Volume size, Volume #) | Testing dataset | Testing data size (Volume size, Volume #) |
|---|---|---|---|---|---|
| Deconvolution with simulated data | **Supplementary Figs. 3, 6** | Phantom objects consisting of dots, solid spheres, and ellipsoidal surfaces. Blurring PSF: 0.8/0.8 NA diSPIM | 128 x 128 x 128<br>100 pairs | Same as training dataset | 128 x 128 x 128<br>9 |
| | **Supplementary Figs. 7** | Simulated spherical beads | 128 x 128 x 128<br>100 pairs | Same as training dataset | 256 x 256 x 256<br>3 |
| Generalization with simulated data | **Fig. 1d**<br>**Supplementary Figs. 5, 7** | Phantom objects consisting of dots, solid spheres, and ellipsoidal surfaces. Blurring PSF: 0.8/0.8 NA diSPIM | 128 x 128 x 128,<br>100 pairs | Simulated spherical beads | 256 x 256 x 256<br>3 |
| | **Fig. 1e**<br>**Supplementary Fig. 4** | | | Human brain phantom | 256 x 256 x 128<br>1 |
| Deconvolution in biological images | **Fig. 2a-c** | U2OS cell, mitochondria, acquired with 0.8/0.8 NA diSPIM | 349 x 512 x 90<br>12 pairs | Same as training dataset | 349 x 512 x 90<br>50 |
| | **Supplementary Video 1** | U2OS cell, mitochondria, acquired with 0.8/0.8 NA diSPIM | 349 x 512 x 90<br>12 pairs | | 284 x 382 x 100<br>200 |
| | **Fig. 2d-e** | Mouse brain neurites acquired by cleared' tissue 0.4/0.4 NA diSPIM | 128 x 128 x 128<br>12 pairs | | 228 x 228 x 1218<br>25 |
| | **Fig. 2g-i,**<br>**Supplementary Fig. 8b,c**<br>**Supplementary Video 2** | Cleared brain tissue slab expressing tdTomato in axons, acquired with 0.7/0.7 NA cleared tissue diSPIM | 256 x 256 x 256<br>40 pairs | | 1500 x 1500 x 42<br>900 |
| | **Fig. 1b, Supplementary Fig. 10c-f, 11a** | *C. elegans* membrane acquired by 0.8/0.8 NA diSPIM | 290 x 364 x 277<br>12 pairs | | 290 x 364 x 277<br>9 |

| | | | | | |
|---|---|---|---|---|---|
| | **Supplementary Fig. 10a-b, 11b** | *C. elegans* nuclei acquired by 0.8/0.8 NA diSPIM | 240 x 360 x 246<br>12 pairs | | 240 x 360 x 246<br>9 |
| | **Supplementary Video 3** | *C. elegans* nuclei acquired by 0.8/0.8 NA diSPIM | 240 x 360 x 246<br>12 pairs | | 240 x 360 x 246<br>291 |
| | **Fig. 3a-c, Supplementary Figs. 12a-c, 16** | U2OS cell ER, mitochondria, lysosome, and Golgi acquired by iSIM | 512 x 512 x 24<br>120 pairs | | 1902 x 1550 x 20 (ER)<br>1920 x 1550 x 32 (Mito)<br>1920 x 1550 x 14 (Lyso)<br>1920 x 1550 x 20 (Golgi)<br>6 |
| Generalization in biological images | **Fig. 3a-b, Supplementary Figs. 12, 13a-b** | U2OS cell ER acquired by iSIM | 512 x 512 x 24<br>120 pairs | U2OS cell mitochondria, lysosome, and Golgi collected by iSIM | 1920 x 1550 x 32 (Mito)<br>1920 x 1550 x 14 (Lyso)<br>1920 x 1550 x 20 (Golgi)<br>6 |
| | **Fig. 3c** | U2OS cell mitochondria acquired by iSIM | 512 x 512 x 24<br>120 pairs | U2OS cell ER collected by iSIM | 1920 x 1550 x 20 (ER)<br>6 |
| | **Supplementary Fig. 13d-e** | Phantom objects consisting of dots, solid spheres, and ellipsoidal surfaces. Blurring PSF: iSIM PSF | 128 x 128 x 128<br>100 pairs | U2OS cell lysosome acquired by iSIM | 1920 x 1550 x 14<br>6 |
| | **Fig. 3e** | Phantom objects consisting of dots, solid spheres, and ellipsoidal surfaces. Blurring PSF: iSIM PSF | | U2OS cell ER collected by iSIM | 1920 x 1550 x 20<br>6 |
| | **Fig. 4a-c Supplementary Fig.14** | Phantom objects consisting of dots, solid spheres, and ellipsoidal surfaces. Blurring PSF: widefield PSF (60X, NA=1.42 oil immerse) | | Fixed U2OS cell acquired by widefield microscopy | 512 x512 x37<br>4 |
| | **Fig. 4d-f** | Phantom objects consisting of dots, | | Widefield *C.* elegans embryos | 1200 x 1200 x 201 |

| | | | | | |
|---|---|---|---|---|---|
| | **Supplementary Fig.15** | solid spheres, and ellipsoidal surfaces. Blurring PSF: widefield PSF (100X, NA = 1.35 silicon oil lens) | | acquired by widefield microscopy | 6 |
| | **Supplementary Fig. 11b** | Phantom objects consisting of dots, solid spheres, and ellipsoidal surfaces. Blurring PSF: 0.8/0.8 NA diSPIM | | *C. elegans* nuclei imaged with 0.8/0.8 NA diSPIM | 240 x 360 x 246<br>1 |
| | **Supplementary Fig. 11a** | | | *C. elegans* membrane imaged with 0.8/0.8 NA diSPIM | 290 x 364 x 277<br>1 |

**Supplementary Table 2, SSIM and PSNR values (means +/- standard deviations from N measurements) comparing RLN, CARE, RCAN and DDN.** RLN always provides the best performance, i.e., highest SSIM and PSNR.

| Figure | Sample | | RLN | CARE | RCAN | DDN | N |
|---|---|---|---|---|---|---|---|
| **Fig. 1d** | Simulated spherical beads | SSIM | 0.97 ± 0.01 | 0.92 ± 0.02 | 0.92 ± 0.03 | 0.93 ± 0.01 | 3 |
| | | PSNR | 35.7 ± 0.65 | 33.5 ± 1.00 | 34.2 ± 0.65 | 34.1 ± 0.72 | 3 |
| **Fig. 1e, Supplementary Fig. 4g** | Brain phantom | SSIM | 0.89 ± 0.01 | 0.86 ± 0.02 | 0.86 ± 0.02 | 0.88 ± 0.02 | 3 |
| | | PSNR | 24.4 ± 0.74 | 22.6 ± 0.40 | 20.0 ± 0.23 | 21.4 ± 0.29 | 3 |
| **Fig. 2b-c** | U2OS cells, mitochondrial label, collected with diSPIM | SSIM | 0.78 ± 0.05 | 0.75 ± 0.04 | 0.76 ± 0.07 | 0.74 ± 0.04 | 3 |
| | | PSNR | 24.1 ± 2.00 | 22.8 ± 1.65 | 23.0 ± 2.23 | 22.3 ± 1.35 | 3 |
| **Fig. 2e** | Mouse brain neurites collected by cleared tissue diSPIM | SSIM | 0.86 ± 0.02 | 0.84 ± 0.03 | 0.76 ± 0.06 | 0.81 ± 0.05 | 3 |
| | | PSNR | 29.7 ± 0.73 | 29.0 ± 1.13 | 23.8 ± 1.54 | 28.3 ± 1.68 | 3 |
| **Fig. 3e, Supplementary Fig. 13f** | U2OS cells, ER label, collected with ISIM, using synthetic mixed structure model | SSIM | 0.69 ± 0.04 | 0.58 ± 0.04 | 0.59 ± 0.05 | 0.61 ± 0.04 | 6 |
| | | PSNR | 22.0 ± 1.76 | 19.9 ± 1.73 | 20.1 ± 1.18 | 20.5 ± 1.42 | 6 |
| **Supplementary Fig. 13c** | U2OS cells, mitochondrial label, collected with ISIM, using model trained on ER | SSIM | 0.97 ± 0.02 | 0.94 ± 0.02 | 0.96 ± 0.02 | 0.95 ± 0.02 | 6 |
| | | PSNR | 40.3 ± 3.43 | 32.1 ± 5.52 | 36.3 ± 4.81 | 35.4 ± 2.91 | 6 |

**Supplementary Table 3, Neural network parameters used in RLN training (RLN-a used the same parameters) and iteration number used in RLD.**

| | Deconvolution ability with simulated data | Generalization ability with simulated data | Deconvolution ability in biological images | | | | | Generalization ability in biological images | | | |
|---|---|---|---|---|---|---|---|---|---|---|---|
| Figure | **Supplementary Figs. 3, 6, 7** | **Figs. 1d, 1e-f Supplementary Figs. 4, 5, 7** | **Fig. 2a-c** | **Fig. 2d, e, g-i, Supplementary Fig. 8b,c** | **Fig. 3a-c, Supplementary Figs. 12a, b, 16** | **Fig. 1b, Supplementary Figs. 10-11** | | **Fig. 3a-c, Supplementary Figs. 12a, b, 13a-c** | **Fig. 3e Supplementary Fig. 13d-f** | **Fig. 4a-c Fig. 4d-f Supplementary Figs. 14, 15** | **Supplementary Fig. 11** |
| Block size | 64 x 64 x 64 | 64 x 64 x 64 | 64 x 64 x 64 | 64 x 64 x 64 | 24 x 64 x 64 | 64 x 64 x 64 | | 24 x 64 x 64 | 64 x 64 x 64 | 64 x 64 x 64 | 64 x 64 x 64 |
| | | | | | | single | dual | | | | |
| Training steps per epoch | 100 | 100 | 120 | 120 | 120 | 120 | 120 | 120 | 100 | 100 | 100 |
| Number of epochs | 200 | 200 | 200 | 300 | 500 | 500 | 500 | 200 | 200 | 500 | 500 |
| Starting learning rate | 0.025 | 0.02 | 0.02 | 0.025 | 0.015 | 0.03 | 0.02 | 0.015 | 0.02 | 0.025 | 0.025 |
| Decay rate | 0.9 | 0.95 | 0.95 | 0.97 | 0.98 | 0.9 | 0.95 | 0.95 | 0.95 | 0.95 | 0.95 |
| Decay step | 200 | 500 | 500 | 250 | 500 | 500 | 500 | 200 | 500 | 1000 | 500 |
| Training time | 2-3h | 2-3h | 2-3h | 3-4h | 3-4h | 4-5h | 4-5h | 2-3h | 2-3h | 3-4h | 3-4h |
| RLD iteration # | Mixed structure:40 | Beads:10 brain:20 | / | 1 (unmatched back projector) | 40 | Membrane :5 Nuclei:10 | / | / | / | 100 | Membrane:5 Nuclei:10 |

**Supplementary Table 4, Parameters used in the training of CARE, RCAN and DDN neural networks.**

| | Generalization ability on simulated data | | | Deconvolution ability on biological samples | | | | Generalization ability on biological samples | | | | | |
|---|---|---|---|---|---|---|---|---|---|---|---|---|---|
| Figure | **Fig, 1d, Fig. 1f, Supplementary Fig. 4g** | | | **Fig. 2b, c, e** | | | **Supplementary Fig. 16** | **Fig. 3e, Supplementary Fig. 13c** | | | **Supplementary Fig. 13f** | | |
| Network | CARE | RCAN | DDN | CARE | RCAN | DDN | RCAN | CARE | RCAN | DDN | CARE | RCAN | DDN |
| Training steps per epoch | 100 | 100 | 100 | 120 | 120 | 120 | 120 | 100 | 100 | 100 | 120 | 120 | 120 |
| Epoch number | 250 | 200 | 200 | 400 | 200 | 200 | 400 | 400 | 200 | 200 | 200 | 200 | 200 |
| Learning rate | 0.0004 | 0.0004 | 0.004 | 0.0004 | 0.0004 | 0.004 | 0.0004 | 0.0004 | 0.0004 | 0.002 | 0.0004 | 0.0004 | 0.002 |
| Decay rate | 0.9 | / | 0.985 | 0.9 | / | 0.985 | / | 0.9 | / | 0.985 | 0.9 | / | 0.985 |
| Decay step | 500 | / | 600 | 500 | / | 600 | / | 500 | / | 600 | 500 | / | 480 |
| Training time | 2-3h | 6-7h | 2-3h | 3-4h | 6-7h | 2-3h | 12-13h | 3-4h | 6-7h | 2-3h | 2-3h | 6-7h | 2-3h |

**Supplementary Video 1:** Time lapse imaging of live U2OS cell transfected with mEmerald-Tomm20, imaged with diSPIM. Lateral (top) and axial (bottom) maximum intensity projections are shown, comparing raw data (single view) vs. RLN predictions. Volumes were acquired with diSPIM every 3 s, 200 time points. See also **Fig. 2a-c**.

**Supplementary Video 2:** 3D rendering of cleared brain tissue slab (~1.4 x 2.3 x 0.5 $mm^3$) expressing tdTomato in axons acquired with 0.7/0.7 NA cleared tissue diSPIM, comparing raw single view, dual-view joint deconvolution and RLN prediction. The RLN prediction improves image resolution and contrast relative to the raw input. The joint deconvolution output causes artifacts and shows many fewer neurites relative to the raw input and RLN prediction, likely due to failures of registration between the two raw views. See also **Fig. 2g-i** and **Supplementary Fig. 8**.

**Supplementary Video 3:** Nuclear imaging (H2B-GFP) in live *C. elegans* embryos. Time-lapse lateral (top) and axial (bottom) maximum intensity projections, comparing raw single view, dual-view joint deconvolution ground truth, single-input RLN and dual-input RLN. hpf: hours post fertilization. See also **Supplementary Fig. 10**.
